# Patterns of diversity under loose genetic draft

**DOI:** 10.1101/2023.07.19.549703

**Authors:** G Achaz, E Schertzer

## Abstract

Recurrent selective sweeps reduce diversity at linked neutral loci, a regime known as genetic draft. Most theoretical work has focused on the tight draft regime, where the selected and neutral loci are closely linked, leading to Multiple Merger Coalescents and a diversity largely insensitive to population size. Here, we investigate the neglected regime of loose genetic draft, where sweeps at a distant linked locus have individually negligible effects but collectively drive diversity. To explore this regime systematically, we make extensive use of the RIF model (Random Initial and Final conditions), a semi-deterministic approximation of selective sweeps that is 10^3^ times faster than standard Wright-Fisher simulations and equally accurate. Using this framework, we derive novel analytical approximations for the coalescence probability under a single sweep, valid for a wide range of recombination-to-selection ratios *A* = *c/s*, and show that they outperform several previous approximations from the literature, which are only accurate for small *A*. Under recurrent loose draft (0.1 *< A <* 0.5), the effective population size scales as a power law of census size, *N*_*e*_ ∝ *N* ^2*A*^, which could contribute to the observed non-linear dependence of diversity on population size. Crucially, despite this strong reduction in diversity, the genealogy converges to a Kingman coalescent, making loose draft patterns indistinguishable from neutrality by standard tests.

## 2 Introduction

The ever-changing environment, where organisms live in, triggers an interrupted sequence of accumulation of newly adaptive traits. As illustrated by the red-queen literary metaphor (Van valen, 1973), one can picture that only organisms that constantly adapt to their environment survive, while the others are doomed to extinction. As most transmitted traits have a genetic component lying in chromosomes, the sequence of successive steps of an adaptive walk is expected to imprint the genome with detectable patterns. In the simplest population model, new beneficial alleles (that correspond to new adapted phenotypes with a relative fitness of 1 + *s*) appear by mutations in the chromosomes of random individuals and rapidly increase in frequency (“sweep”) up to their complete fixation. Just after the completion of a selective sweep, every individual bears the beneficial allele at the selected locus and the diversity at this locus is entirely wiped out. In this sense, it can be thought that positive Darwinian selection purges out diversity extremely efficiently.

To acknowledge the observation that higher than expected genetic diversity is segregating at most loci for most species, it has been proposed that selective sweeps were both rare at the population genetics time scale and sparse along the genome. More precisely, under the neutral theory of molecular evolution (Kimura, 1968; King and Jukes, 1969; Kimura, 1983), genetic variations segregating at observable frequencies are assumed to be neutral and, on first order approximation, free from selection. In the neutral framework, the fate of neutral variants is entirely driven by genetic drift, which purges diversity much less efficiently than selection. Indeed, as the effect of genetic drift is inversely proportional to the population size *N*, the loss of diversity occurs at the scale of *N* generations (Wright, 1931). Although these derivations are directly applicable to a theoretical Standard Neutral Model, named the Wright-Fisher model (Ishida and Rosales, 2020), one usually replaces the census size *N* by an effective population size *N*_*e*_ to characterize the effect of genetic drift in other models or in biological data. A long list of natural add-ons to the simple haploid Wright-Fisher model can result in (important) differences between *N* and *N*_*e*_ (Charlesworth, 2009; Waples, 2022). Typically, extra features include: large demographic variations, recent foundation time of the species following recurrent species fragmentation, strong structuration in local demes, sex ratios, large variance of offspring number, etc. Shortly after the neutral theory was established, Lewontin (1974) observed that molecular diversity (and thus 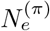, estimated from it) varies across species within a much narrower range than population size does, suggesting that census population size *N* is a weak predictor of genetic diversity. This discrepancy between *N* and 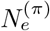 has since become known as Lewontin’s paradox of variation (Leffler *etal*., 2012; Ellegren and Galtier, 2016; Galtier and Rousselle, 2020; Charlesworth and Jensen, 2022).

In this article, we analyze the interplay between selection and drift, studying the fate of beneficial alleles in a finite Wright-Fisher haploid population. As a rapid survey, one can compare the time scale of the fixation of neutral alleles by genetic drift, *O*(*N*), and the fixation of beneficial alleles by selective sweeps, *O*(1*/s*). This immediately suggests that when *Ns* ≪ 1, the time scale of selection is very long compared to the drift time scale, and in practice the random sampling of genetic drift will dominate the system and will drive entirely the fate of the allele. On the other end, when selection is strong, selection acts as a dominating evolutionary process. Yet, even in a strong selection regime (*Ns* ≫ 1), the effect of genetic drift cannot be neglected when the beneficial allele is at low (or high) frequency and when selection operates at the scale of 1*/N* ≪ *s* ≪ 1. As a direct consequence, in this parameter range, the probability of fixation of a new beneficial allele is well approximated by 2*s* (Haldane, 1927), illustrating that most of the time, beneficial alleles are lost by genetic drift.

Without recombination, a single beneficial mutation completely drives the fate of the entire chromosome, provided it is not lost by genetic drift. This effect of linkage is central in mitochondrial chromosomes that lack recombination (Hill, 2020).

A richer view of the interplay between selection and drift explicitly models the genome as a sequence of loci between which recombination can take place. The influence of a nearby selected allele on a focal neutral locus has been named *linked selection*. Both positive linked selection, named hitchhiking (Maynard smith and haigh, 1974), and negative linked selection, named background selection (Charlesworth *et al*., 1993), reduce diversity in surrounding loci. The strongest empirical support for the reduction of diversity through linked selection stems from the observation that regions with low recombination show less diversity than regions with high recombination (AGUADE *etal*., 1989; Stephan and Langley, 1989; Begun and Aquadro, 1992). Indeed, both positive and negative regimes of linked selection reduce drastically surrounding diversity for tightly linked loci. Interestingly, this effect is stronger for abundant species (Corbett-Detig *etal*., 2015), suggesting that for large *N*, linked selection is more intense, as is selection more generally.

In the hitchhiking model, a beneficial mutant of selective coefficient *s* appears at a selected locus located at a (genetic) map distance *c* of a neutral locus. Just after the end of the selective sweep, the ancestral diversity at the selected locus has been entirely wiped out, as all individuals are descendants of the ancestral genome where the favorable mutation occurred. On the contrary, on the neutral locus, some diversity is preserved, as the beneficial allele during its rapid expansion is being recombined to other genomes, thus preventing them to be all clonal. The further away from the selected locus, the more recombination and the less the reduction of diversity. The conclusion of Maynard smith and Haigh (1974) is that diversity at the neutral locus is strongly lowered by the selective sweep when the coefficient *A* = *c/s* is small: selection is strong (*s* large), recombination is low (*c* → 0) or both. When *A* = 0, there is no recombination during the sweep and diversity at the neutral locus equals diversity at the selected locus. For (very) strong sweeps all diversity is removed from the region. At the opposite, when *A* ≫ 1, the two loci are completely independent and the neutral locus is unaffected by the selective sweep. Although the original derivation of the model assumed an infinite population size and was mostly focused on the reduction of heterozygosity, subsequent works refined the results and eventually rephrased it in a diffusion setting (Wiehe AND Stephan, 1993) or in a coalescent framework (Kaplan *etal*., 1989; Fay and Wu, 2000). The hitchhiking effect has further been connected to the effect of inflation of the additive variance in fitness (Barton, 2000), originally reported by Robertson (Robertson, 1961). Broader scenarios of hitchhiking effects were studied in a fluctuating environment, spatial structure and soft sweeps (Barton, 2000; Hermisson and Pennings, 2005; Pennings and Hermisson, 2006).

To model the constant adaptation of organisms, one can envision a constant flow of beneficial mutations at the loci under selection. As the effect of recurrent hitchhiking on the neutral locus mimics (partially) some features of genetic drift (e.g. fluctuation of the frequencies of the alleles at the neutral loci), the regime has been baptized genetic draft (Gillespie, 2000A,b). In this regime, it is the repeated occurrence of beneficial mutations at the selected locus that dictates the fate of neutral alleles and conversely the coalescence process at the neutral locus. Although the original derivation was done for two lineages and infinite population size (Gillespie, 2000A,b), several approximations were derived for finite population and *n* lineages (Durrett and Schweinsberg, 2004; Etheridge *etal*., 2006; Coop and Ralph, 2012). As this effect of hitchhiking decreases with *A*, most of the previous work has focused on the strong selection, low-recombination regimes (*A* → 0), with a notable exception of Barton (1998). At the limit, the genealogy of a sample of *n* individuals is well described by a Multiple Merger Coalescent (MMC) process (Pitman, 1999; Sagitov, 1999; Tellier and Lemaire, 2014; Berestycki, 2009). In MMC, several lineages (the ones that are captured by the selective sweep) coalesce almost simultaneously once the time has been properly rescaled, assuming the time of a sweep is much smaller than the time elapsed between two consecutive sweeps. Although the probability of coalescence in most approximations (Wiehe and Stephan, 1993; Barton, 1998, 2000; Durrett and Schweinsberg, 2004; Etheridge *etal*., 2006) is a power law of *N* (*P*_coal_ ∝ *N* ^−2*A*^), it becomes independent of *N* at the limit of *A* → 0. This has led to a view where the coalescent process is entirely driven by the frequency of the selective sweeps and does not depend anymore on the population size (Coop and Ralph, 2012).

In this article, we show that contrarily to the regime *A* ≪ 1 alluded to in the previous paragraph, larger values of *A* generate another regime that we call *loose genetic draft*, whose patterns of diversity (e.g. pairwise difference and Site Frequency Spectrum) are almost indistinguishable from neutrality, with the caveat that this quasi-neutrality emerges on a time scale smaller than the time scale of the neutral model (*N* ^2*A*^ vs *N*). We name *tight hitchhiking* the regime where recombination is relatively weak compared to selection, characterized by a small recombination-to-selection ratio (1*/N* ≪ *s* ≪ 1 and *A* ≪ 1); the hitchhiking is tight in the sense that fixation of the advantageous selected allele has a sizeable effect on the genetic diversity at the tightly linked neutral locus. After completion of the sweep, a positive fraction of the population inherits the genetic material associated to the mutant type at the neutral locus. In the absence of new mutation at the neutral locus, the heterozygosity is entirely lost after one or few sweeps. In contrast with the aforementioned works, we study here the *loose hitchhiking* regime (1*/N* ≪ *s* ≪ 1 and *A* = *O*(1)), where recombination is high enough so that, at first approximation, the evolution of the genetic diversity at the neutral and the selected sites are almost independent along the course of a single sweep. Although the effect of a single sweep is negligible in this regime, we show that when sweeps are repeated at a high enough frequency (more frequent than the drift scale), their infinitesimal effects accumulate and have a sizeable aggregated impact — larger than standard genetic drift — for large populations *N* → ∞.

A key challenge in exploring this regime systematically is computational: accurately simulating selective sweeps in finite populations across a wide range of parameters is expensive in CPU time as Wright-Fisher forward simulations scale quadratically with population size (*N* generations of *N* individuals). To overcome this, we make extensive use of the *RIF model* (Random Initial and Final conditions), a semi-deterministic approximation of the fixation trajectory, previously described by Martin AND Lambert (2015). The core idea of the RIF model is that a selective sweep in a large finite population is almost entirely deterministic at intermediate allele frequencies, while stochasticity is confined to two brief phases: the initial spread from low frequency and the final approach to fixation. Both phases are well approximated by branching processes, and by Yaglom’s law each converges to a deterministic exponential trajectory seeded by a random initial condition drawn from an exponential distribution with mean 1*/*(2*Ns*). The trajectory between these two random endpoints — the initial frequency *x*_0_ ∼ Exp(2*Ns*) and the final frequency *x*_*τ*_ ∼ 1 − Exp(2*Ns*) — is then fully deterministic, following a logistic equation. This seemingly simple approximation turns out to be remarkably accurate: it matches Wright-Fisher simulations in both forward properties (allele frequency distributions, fixation times) and backward properties (pairwise coalescence probabilities) across the full parameter range we explore. Crucially, it is also dramatically more efficient: a single RIF realisation requires drawing just two exponential random variables, making it approximately 10^3^ times faster than a naive Wright-Fisher simulation (0.3 seconds vs. 4 minutes for 10^5^ replicates at *N* = 10^5^, *s* = 0.01). This computational efficiency is not a minor convenience: it is what makes it tractable to explore the large parameter space of loose genetic draft, spanning population sizes from 10^2^ to 10^8^ and *A* values from 0.01 to 0.5, that would otherwise be prohibitively slow. In this sense, the RIF model is the methodological backbone of this study that, we believe, could be used to explore more complex biological scenarios involving selection and recombination.

In this article, we first provide strong evidence that a selective sweep in finite population is well approximated by the RIF model, and establish its forward and backward properties. We provide analytical approximations when population size is very large and compare them to preexisting approximations from the literature, discussing their strengths and weaknesses. Then, using the RIF model, we explore patterns of diversity under the regime of genetic draft, especially focusing on the neglected loose genetic draft regime. In particular, we show that for large enough *N* and *A* values (large population–loose hitchhiking), the genealogical structure at the neutral locus becomes indistinguishable from a neutral genealogy. More formally, the coalescent tree converges to a Kingman tree (neutral SFS), except for its reduced time scale (small *π*). We end by discussing the parameter space on which the model holds and how it fits observed patterns of molecular diversity.

## 3 Model derivation and Results

### 3.1 The RIF model: logistic equation with Random Initial and Final frequencies

#### 3.1.1 Selective sweep as a diffusion

In order to motivate the RIF model, we build our approach on a diffusion approximation of a haploid population of constant size *N*, in which a derived allele at frequency 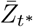 benefits from a relative fitness (1 + *s*) compared to the ancestral alleles of relative fitness 1. The trajectory of this beneficial derived allele is modeled by a Wright-Fisher diffusion with selection, provided that usual diffusion conditions hold (*N* ≫ 1 and *s* ≪ 1), for which of one time unit of *t*^∗^ corresponds to 1 discrete Wright-Fisher generation:

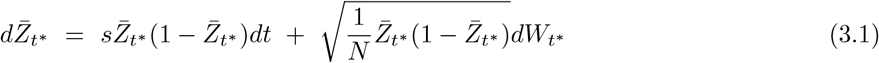

where *W* is a standard Brownian motion. To model a fixation trajectory, we condition the latter diffusion on fixation and denote by *Z* the resulting process. To construct the conditioned process, we proceed in two steps (1) we start from 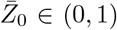 and we follow the frequency of a fixing allele by first conditioning (3.1) on survival, and (2) we let the initial frequency go to 0 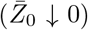. Note that although the initial frequency of a Wright-Fisher model is 1*/N*, the resulting diffusion approximation starts from 0 for a large population. The resulting conditioned process *Z* can be shown to be itself a diffusion obtained from the unconditioned diffusion through a Doob *h*-transform (Etheridge *et al*., 2006):

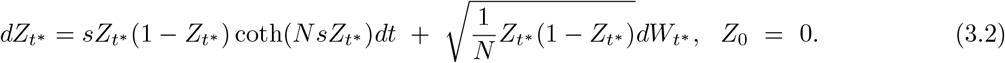

(Note that we remove the bar to indicate the conditioning process.) The effect of the conditioning is to multiply the first term by coth(*NsZ*_*t*_∗). Since

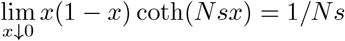

*Z* is repulsed away from 0 at small frequency. The process thus starts from 0 and is repulsed away from the origin so that it never returns to 0 on later times. ^∗^

From now on, we will assume a selection regime where

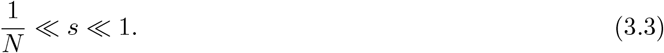

Under this regime, it is convenient to accelerate time by a factor 1*/s* and thus introduce a rescaled process *z*_*t*_ := *Z*_*t*_∗ _*/s*_. In this new time scale *t* represents 1*/s* units of *t*^∗^ (that were Wright-Fisher generations). A direct computation (dividing both sides of eq. (3.2) by *s*) yields to:

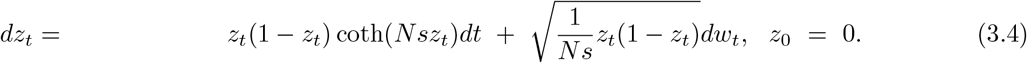

where 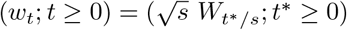 is distributed as a standard Brownian motion.

#### 3.1.2 From selective sweep to RIF

The aim of the present section is to show that the diffusive selective sweep (3.4) is well approximated by a deterministic logistic equation where the initial frequency and the final frequency are characterized by independent exponential random variables with mean 1*/*(2*Ns*). We name this approximation of the fixation trajectory the RIF-model, for Random Initial and Final conditions. This model has been previously described in the literature (Martin and Lambert, 2015) under the name of semi-deterministic. See also (Desai and Fisher, 2007; Charlesworth, 2020; Barton, 1998) for a similar approach. In their work, the authors derive this approximation starting from a Moran-like model. In this manuscript, we provide rapid insights on its justification in the diffusion setting.

The idea of the RIF model relies on the classical observation that the fixation trajectory is almost deterministic at intermediary frequency, and that this deterministic regime is interpolated by two stochastic phases – respectively at low and high frequency of the mutant allele – which are well approximated by branching processes.

Let us now spell out the argument in more details. When *z*_*t*_ is at intermediate frequency, since *Ns* ≫ 1 (3.4) is well approximated by the deterministic logistic equation (time *t* is still in 1*/s* generations):

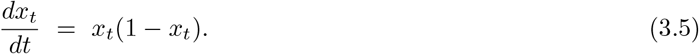

To fully determine the deterministic dynamics, it now remains to determine the initial condition of the system. The idea is to observe the process at small frequency (i.e., when *t* = *o*(1)) and to observe that under (3.5)

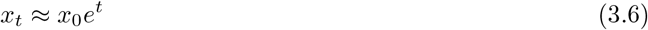

so that it remains to identify *x*_0_ such that the latter approximation holds at small times for the diffusion equation.

Set *y*_*t*_ := *Nsz*_*t*_. At low frequency (*i*.*e*. at small time *t* such that *z*_*t*_ ≈ 0), (3.4) writes

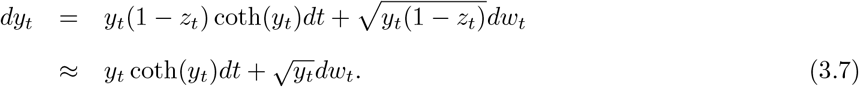

The latter approximation is identical in law to a supercritical continuous-state branching process (Feller diffusion, see e.g. (Lambert, 2008)) conditioned on eternal survival ^†^. In the fixed size Wright-Fisher populations with directional selection, this Feller diffusion approximation translates into approximating the advantageous alleles at low frequency by a supercritical branching process (a classical approach used by Fisher and Haldane, see review by Patwa and Wahl (2008)) conditioned on survival.

A key property (known as the Yaglom’s law) of the process *y* (described by the approximation (3.8)) is that it converges to a deterministic exponential growth with a random initial condition

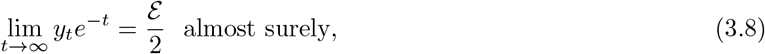

where ℰ is a standard exponential random variable (of mean 1). Since 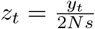, we can informally write:

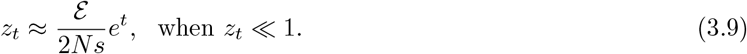

Comparing this estimate with (3.6), we conclude that the process *z* observed at intermediary frequency is well approximated by the deterministic solution *x* started from the initial condition

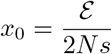

Finally, it has been noted by Coop and Griffiths (2004) that the Wright-Fisher fixation trajectory *z* is reversible in the sense that if *τ* denotes the fixation time then

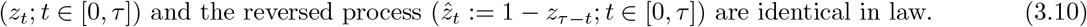

By time reversal, this shows that the fixation time is well approximated by stopping the deterministic trajectory at

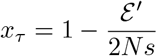

where ℰ′ is an independent standard exponential random variable.

To summarize, the previous heuristics show that one can approximate the initial stochastic phase by an instantaneous jump from frequency 0 to *x*_0_ and the final stochastic phase by an instantaneous jump from frequency *x*_*τ*_ to 1. The two phases are interpolated by a deterministic logistic phase. In the following, this approximation will be referred to as the RIF model.

### 3.2 Properties and validity of the RIF model

In this section, we will list several properties of the RIF model. To assess the validity of the RIF model, we will compared individual based Wright-Fisher simulations of trajectories conditioned on fixation (using a naive rejection algorithm) with forward RIF-model realisations with random *z*_0_ and *z*_*τ*_ (sampled by Monte-Carlo).

#### 3.2.1 Forward perspective

We compute two distributions. First, we report the distribution of allele frequency at the rescaled time *t* = ln(*Ns*), where the trajectory has become deterministic (Figure 1, left panel). Second, we report the distribution of the fixation time (Figure 1, right panel). In the RIF model, the rescaled fixation time of the trajectory is given by:

**Figure 1.**
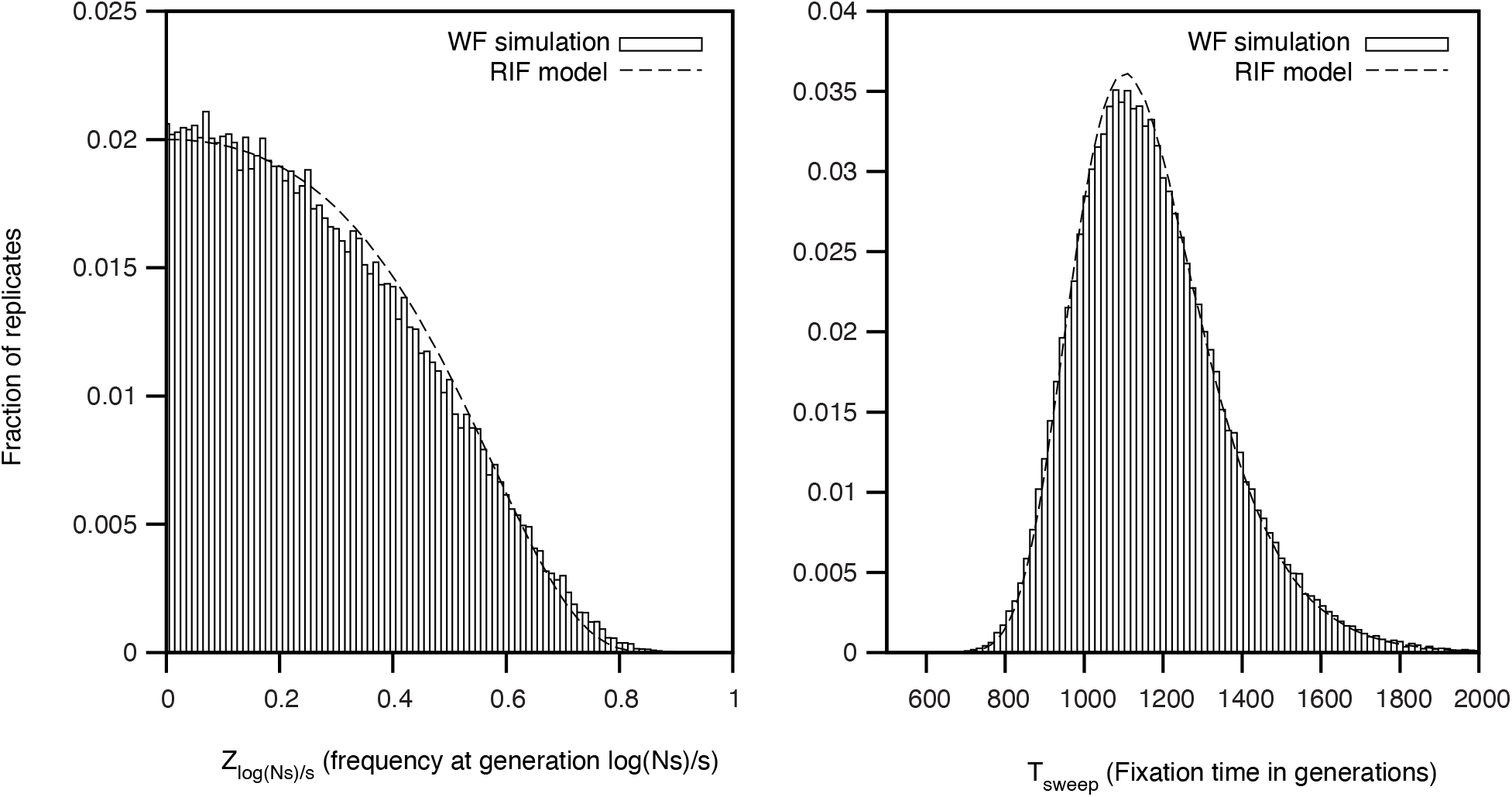
We compare the outputs of simulated selective sweeps in a Wright-Fisher model (discarding replicates where the beneficial allele is lost) to the prediction of the RIF model. Population size is *N* = 10^5^ and the coefficient of selection is *s* = 0.01. On the left panel, we display the distribution (bins of 0.01) of frequency *Z*_*t*_∗ of the fixing beneficial allele at the generation *t*^∗^ = ln(*Ns*)*/s*. The RIF-model corresponds to eq. (3.9), with ℰ generated by Monte-Carlo. On the right panel, we display the distribution (bins of 15 generations) of fixation time. The RIF-model corresponds to equation 3.11, with ℰ and ℰ′ generated by Monte-Carlo.

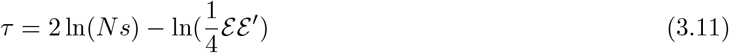

implying that the mean fixation time in generations is well approximated by

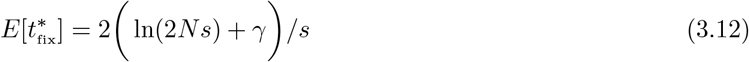

where *γ* is the Euler constant (*γ* ≈ 0.5772).

Figure 1 shows that both distributions (derived allele frequency at time ln(*Ns*)*/s* and fixation time) display an excellent numerical agreement between both the Wright-Fisher simulations and the RIF-model realisations. They illustrate how the RIF-model constitutes an excellent approximation for selective sweeps in finite populations. Diffusion realisations based on eq. 3.2 also display excellent numerical agreement (data not shown).

##### Numerical simulations

A single numerical realisation of the RIF model is instantaneous as it only relies on randomly drawing two exponential random variables (*z*_0_ and *z*_*t*_), whereas a forward realisation of a sweeping allele in a Wright-Fisher framework (only following frequencies) requires as many binomial samplings as the fixation time (plus few samplings for the runs where the beneficial allele is lost). To simulate using the diffusion approximation, conditioned on fixation (see eq. (3.2)), one draw as many normal random variables as time increments *dt*. Both Wright-Fisher and diffusion increase CPU time with population size since selective sweeps take more time to complete. As a crude comparison, the realisation of 10^5^ replicates of selective sweeps (*N* = 10^5^ and *s* = 0.01 as in Figure 1) took an a laptop: 0.3” for the RIF model, 2’45” for diffusion with *dt* = 1 and 4’01” for Wright-Fisher with rejections of losses. This makes the RIF approximation 10^3^ times faster than a naive Wright-Fisher version. In this regard, the RIF model can be preferred in *most* numerical simulations of strong selective sweep (*Ns* ≫ 1), both forward or backward (see below).

#### 3.2.2 Backward perspective: coalescence probability under the RIF model

##### Model assumptions

We now turn on describing the backward coalescent process associated to the RIF model, and compare it with the coalescence process associated with a forward selective sweep in finite population. From now on, *x* will refer to the RIF model, whereas *z* will refer to the diffusion model.

As time is now counted backward, we define a reverse trajectory into the past noted 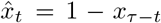. Since time is reversed, 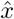 starts from the initial condition 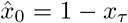, where *x*_*τ*_ = ℰ′*/*2*Ns*. 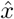 terminates at 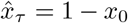, where 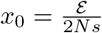. Recall that time is rescaled by 1*/s* (see Section 3.1.1 and eq. 3.4), so that the recombination rate becomes *A* = *c/s*. Given the reversed fixation trajectory 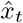, we follow the construction of Hudson and Kaplan (1988) who model the genealogy at the neutral locus as a structured coalescent process. For a schematic view of this structured coalescent is provided in Figure 2.

**Figure 2.**
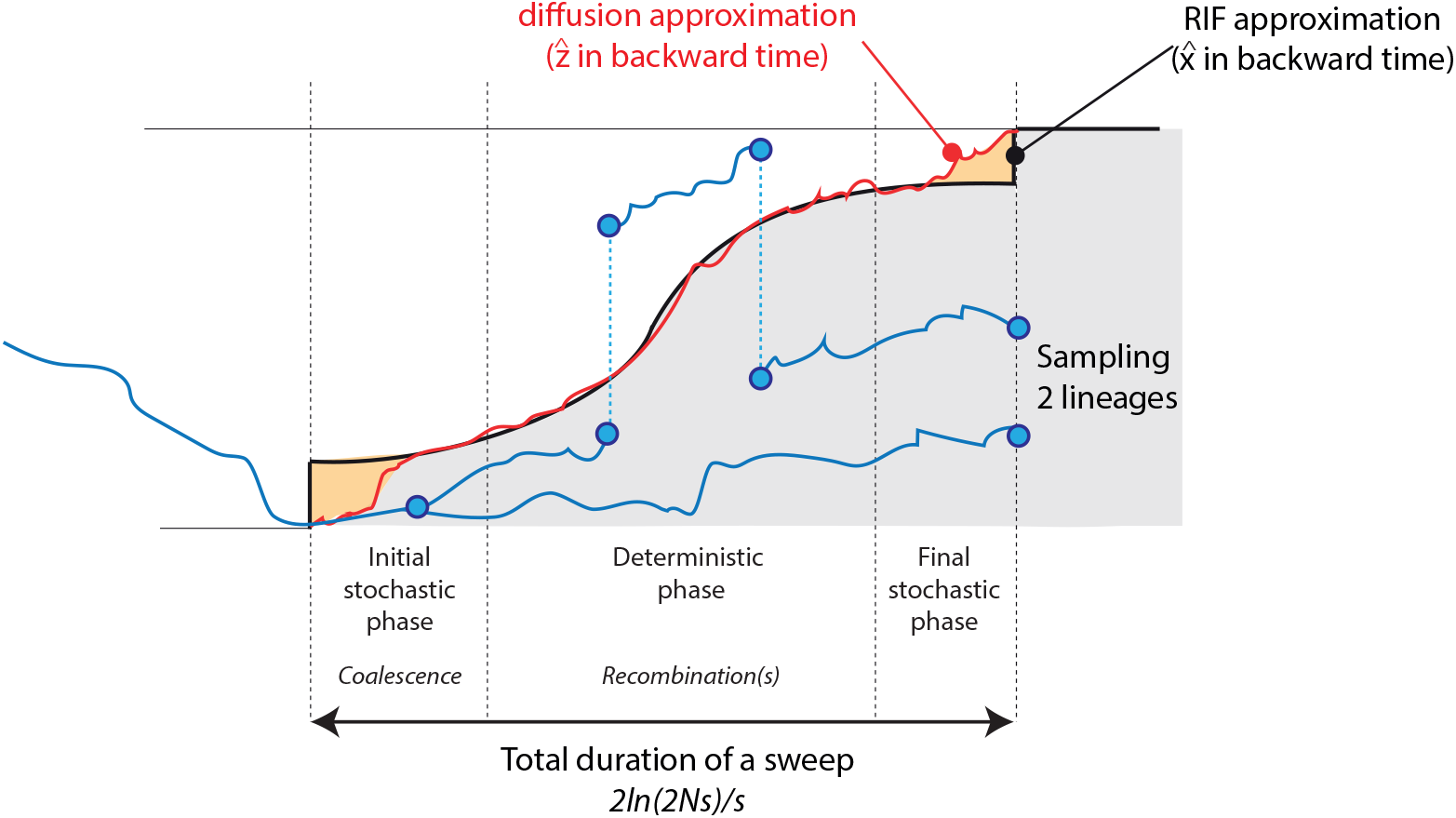
A model of structured coalescent. In this sketch of the model, we show a single selective sweep event that lasts on average ≈ 2 ln(2*Ns*)*/s*. Two lineages at a linked neutral locus (represented in blue) are sampled after the completion of the selective sweep. On their journey backward in time, the lineages start linked to the beneficial allele at the selected locus (they are in the gray area). They can nonetheless recombine back and forth during the selective event changing their association to the ancestral allele or back to the beneficial allele. Although coalescence could in principle occur anytime during the whole sweep, it occurs in practice only at the onset of the selective sweep (as justified in the appendix). We also illustrate the two approximations used to derived the probability of coalescence: the RIF approximation (in black) and a diffusion approximation (in red).

1. Lineages can be of type 0 (linked to the beneficial allele – grey zone on Fig. 2) or type 1 (linked to the ancestral allele). The two lineages start as type 0.
2. Lineages transition from type 0 to type 1 at effective recombination rate 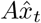 and from type 1 to type 0 at effective rate 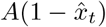.
3. Coalescence can only occur within lineages of the same type: pairs of type 0 coalesce at rate 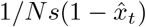 and pairs of type 1 coalesce at rate 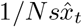. Coalescence rates are multiplied by 1*/s* since time has been rescaled by the same amount.
4. In addition, we also assume that if two lineages are of type 0 at time *τ*, the two lineages trace back to the ancestral mutant and are forced to coalesce.

##### Numerical simulations

We compute the mean coalescence probabilities for two types of Monte-Carlo simulations. The first one is a Wright-Fisher forward-then-backward (like in e.g. msms (Ewing and Hermisson, 2010)), where the fixation trajectory is simulated forward and a structured coalescent is then simulated backward conditioned by this trajectory as in Simonsen *et al*. (1995). The second is a simple structured coalescent assuming a backward RIF-model realization (randomly generating *x*_0_ and *x*_*τ*_), that is the RIF model, without further analytical approximation.

In Figure 3, we report results for either *A* fixed (panel A) or *N* fixed (panel B). We clearly see how the two types of Monte-Carlo simulations are numerically almost indistinguishable, again pointing to the power of using a CPU efficient RIF-model, here for backward coalescent numerical simulations.

**Figure 3.**
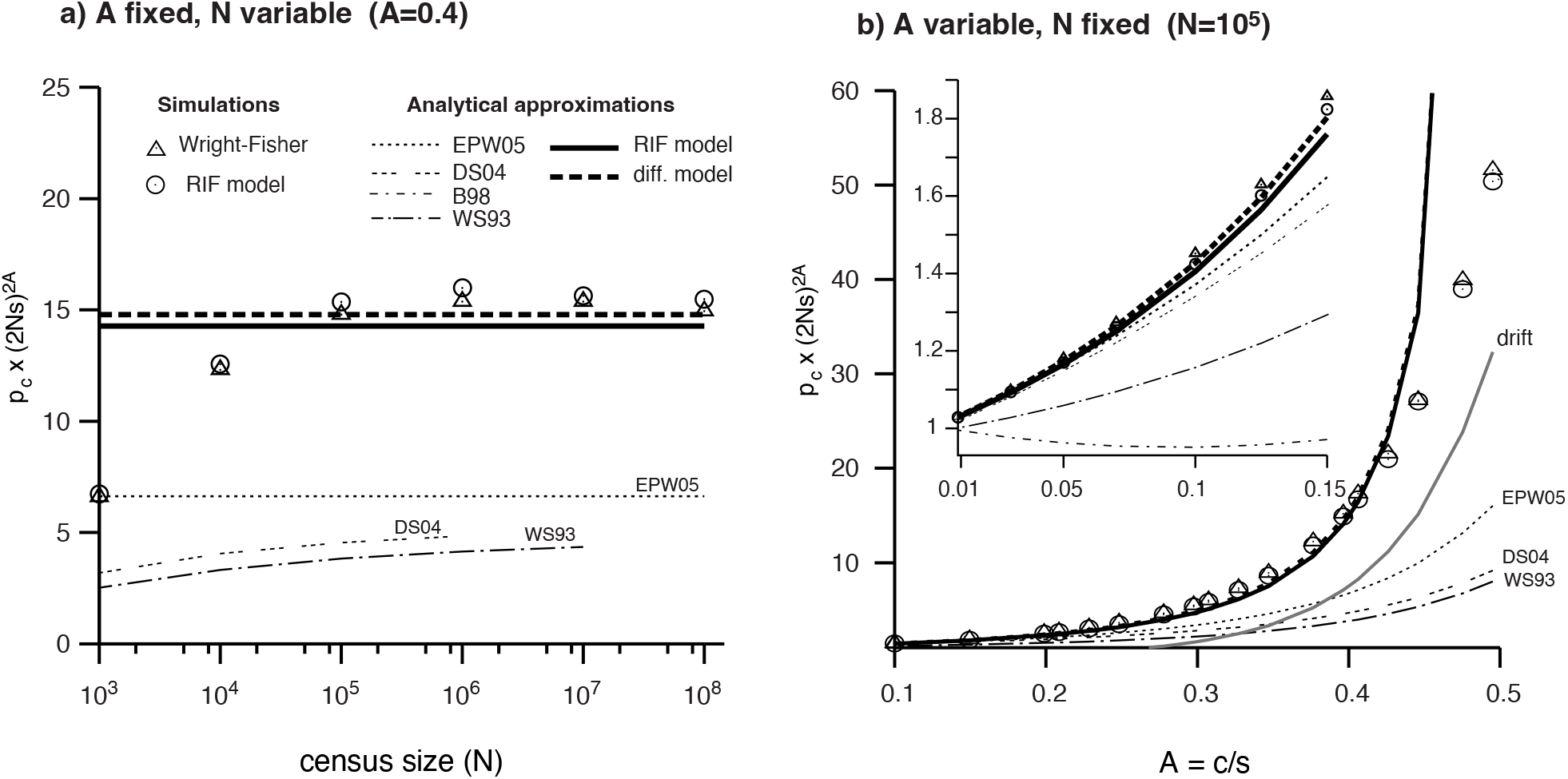
Numerical evaluation of coalescence probability. We compare the outputs of two different simulation procedures with six analytical approximations of *p*_*c*_ × (2*Ns*)^2*A*^ the probability of coalescence deprived of the first term. For all cases, the selection coefficient of the sweeping allele is *s* = 0.01. The ‘Wright-Fisher’ simulations are discrete non-overlapping generations of a Wright-Fisher model with selection ignoring cases where the beneficial allele is lost. The ‘RIF model’ is a deterministic sweep with random initial and final frequencies. WS93, DS04, EPW05 are the analytical approximations provided respectively in Wiehe and Stephan (1993); Durrett and Schweinsberg (2004); Etheridge *et al*. (2006). WS93 and DS04 could not be computed for large *N* . The two other expressions are derived in the present work (‘RIF model’ is eq. (3.14) and ‘diff model’ is eq. (3.15). a) On this panel, the ratio *A* = *c/s* is fixed to 0.4 and the population size varies from 10^3^ to 10^8^. b) On this panel, the population size *N* is fixed to 10^5^ and the *A* ratio varies from 0.01 to 0.5. The insert is a zoom for small *A* values. The “drift” curve is computed using the approximate coalescence probability by drift during the sweep: 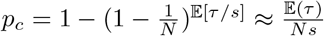.

##### Approximating RIF coalescence probability for large populations

There is no closed-form expression for the mean coalescence probability 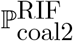 of two lineages under the RIF model. However, as shown in Fig. 3 (panel A), the rescaled quantity

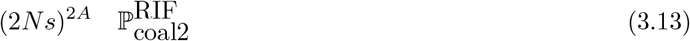

becomes independent of *N* for sufficiently large *N* . In the Appendix (see Section A.1), we justify this numerical observation by showing that as *Ns* ≫ 1, we have

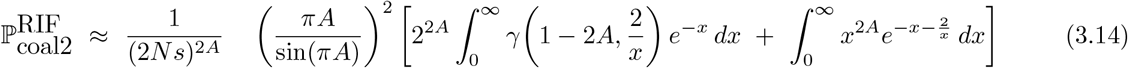

##### RIF approximation concurs with diffusion approximation

Recall that the RIF model was motivated by the diffusion model introduced in Section 3.1.1. How does the latter approximation compare with the mean coalescence probability computed from the diffusion model itself? In the Appendix (see Section A.2), we show that when *Ns* ≫ 1 is large, the mean coalescence probability under the diffusion model can likewise be approximated by

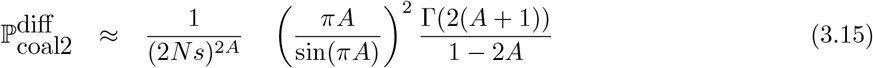

Both approximations (Eqs. (3.14) and (3.15)) can be decomposed into three factors reflecting the selective sweep, as illustrated in Fig. 2. The first factor, (2*Ns*)^−2*A*^, corresponds to the average duration of sweep if we remove the last stochastic phase (≈ ln(2*Ns*)*/s*) during which lineages must not recombine (*e*^−*rτ*^), and sets the global timescale. The remaining two factors arise from a finer description of lineage histories during the sweep. During the “deterministic phase” (see Fig. 2), lineages may recombine, and the second factor — (*πA/*sin(*πA*))^2^ — gives the probability that both lineages are of type 0 at the end of this phase (first vertical dashed line from the left in Fig. 2). The difference between the two approximations reflects the distinct behavior of the two processes during the “initial stochastic phase”, that is, the phase in which the early-time fluctuations of the frequency process *z* play a role. Despite this apparent difference, both approximations yield very similar numerical values (see Figure 3(Panel A)).

We close this section by noting that the diffusion approximation Eq. (3.15) is related to an expression derived by Barton (1998) (their Eq. 13) by other methods, up to a minor modification. In Barton (1998), it is claimed that for a Wright–Fisher model the coalescence probability is given by

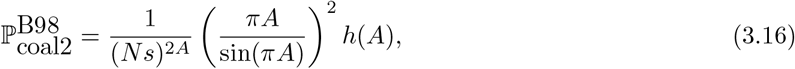

where the function *h* is implicitly defined as follows ^‡^. Consider the ODE

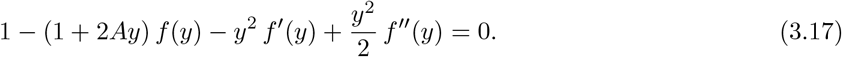

In the words of that author, *h* is determined by requiring that the solution behaves asymptotically as *h*(*A*)*/y*^2*A*^ for large *y*, together with the boundary condition *f* (0) = 1. In the Appendix (see Section A.3), we show that we can indeed recover our formula from the ODE, given some corrections to Barton (1998).

In summary, we find that the approximation derived for the RIF model (Eq.(3.14)) is consistent both with the large-*N* asymptotics of the underlying diffusion model (Eq.(3.14) and with the corrected Barton (1998) Formula (3.16).

##### Comparison with other approximations

We now consider the two expressions derived earlier (under the RIF and the diffusion model) together with previous work of Wiehe AND Stephan (1993),(Durrett and Schweinsberg, 2004) and Etheridge *et al*. (2006)

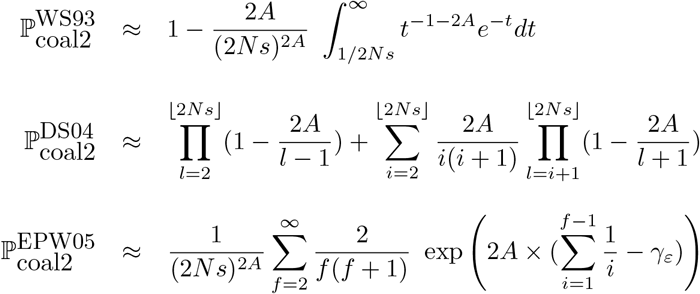

We compare all formulas with the simulations. Although, our two approximations provide very close numerical values, the approximations in Wiehe and Stephan (1993); Durrett and Schweinsberg (2004); Etheridge *et al*. (2006) Are only satisfactory for low *A* values: typically *A <* 0.1. In particular, we see from panel B Figure 3 that all previous approximations predict a coalescence probability lower than the neutral case when *A* is large enough. In addition, our two approximations become better when *N* gets larger (Panel A of Figure 3) and when *A* is not too close to 0.5 (Panel B of Figure 3). Although *A* could take in principle any values in [0, ∞], our approximations only hold when *A <* 0.5, and even more when *A* is not too close to 0.5, where the approximations tend to infinity.

This discrepancy stems from the fact that Wiehe and Stephan (1993), Durrett and Schweinsberg (2004), and Etheridge *et al*. (2006) all assume tight linkage (*A* ≪ 1), under which coalescence events preceded by recombination are negligibly rare and can therefore be ignored. This approximation, however, breaks down when *A* is in between 0.1 and 0.5.

In summary, care must be taken when applying analytical approximations from the literature. Under loose linkage, we find that only a modified version of the result from Barton (1998) yields accurate predictions for sufficiently large *Ns*.

##### RIF approximation vs the RIF model

In Figure 4, we report the probability of coalescence of two lineages as a function of *A* = *c/s* from *A* = 0.01 that corresponds to a tight draft regime to *A* = 5 that is a drift regime.

**Figure 4.**
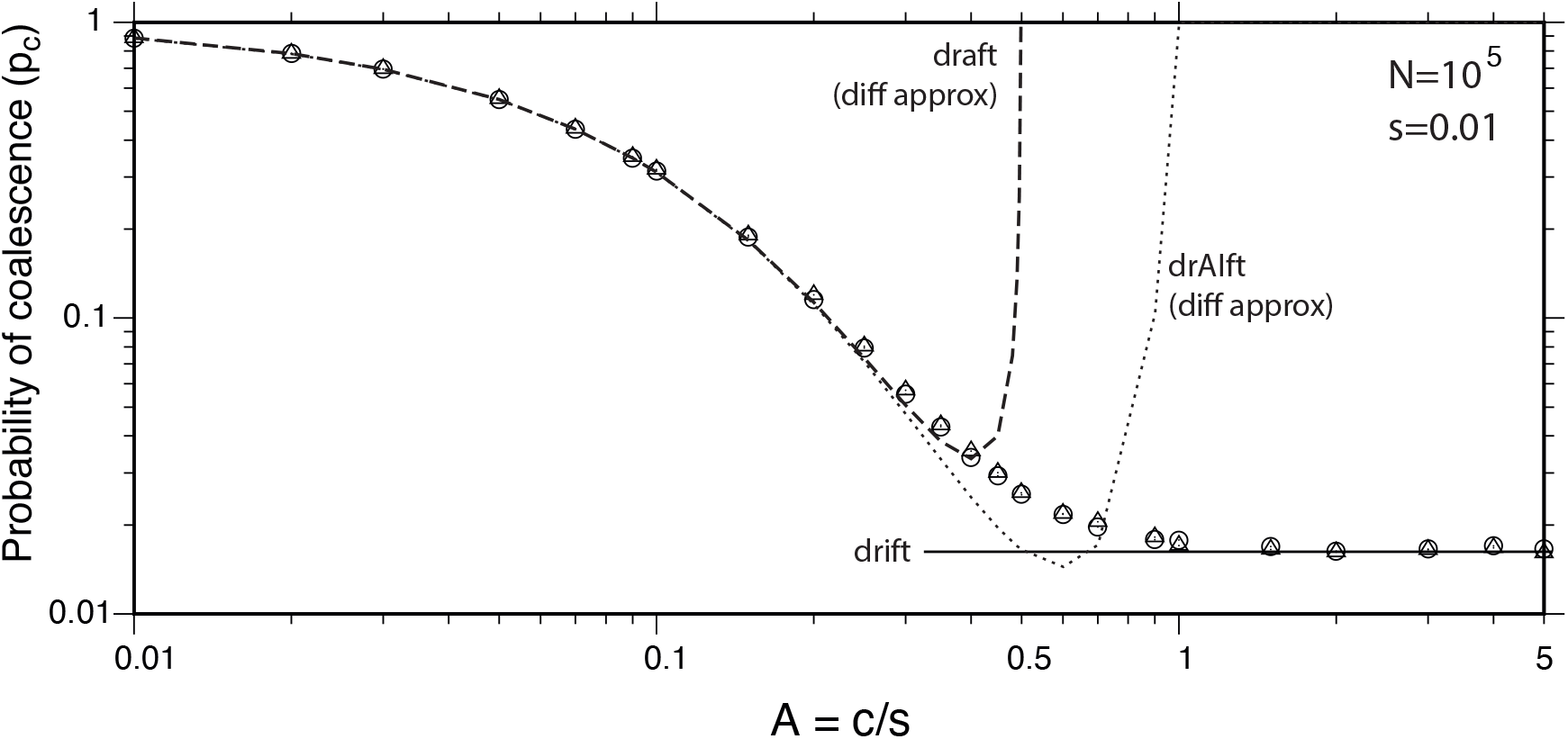
From draft to drift. The probability of coalescence is plotted as a function of the *A* = *c/s* ratio, for a census size of *N* = 10^5^ and a selective coefficient of *s* = 0.01. the dashed line represents the approximation derived from diffusion (eq. 3.15), the dotted lines a second order approximation we named drAIft (eq. A.23) and finally the drift line is 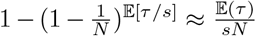. Like in Figure 3, circles are averages of forward Wright-Fisher simulations and triangles from backward RIF simulations (both with 10^6^ replicates).

Results shows that our approximation of the coalescence probability (eq. 3.15) is very good when *A <* 0.4 for a population size of 10^5^ but becomes unreliable when *A* → 0.5, where it is infinite. Extra work would thus be needed to analytically characterize the transition between genetic draft and genetic drift in the critical region surrounding *A* = 0.5.

We also note from Figure 3, that the RIF approximation is only accurate for large populations (*N* ≥ 10^5^) and tends to overestimate the mean coalescence probability contrarily to the exact RIF model that is always in exact agreement with Wright-Fisher individual-based, irrespective of population size or the value of *A* (Figure 4).

##### The drAIft approximation

Although, the approximations are only valid for *A <* 0.4, we derive a second order correction in Appendix A.4, where we admits a finite size correction to the diffusion approximation (eq. (A.23)) that regularizes the divergence at *A* = 1*/*2. In particular, for *A* ∼ 1*/*2, the coalescence probability reduces to:

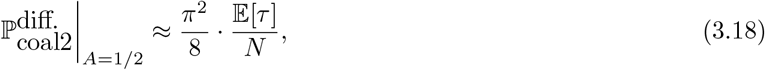

which can be compared to the drift coalescence probability during a sweep, 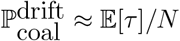. This indicates that at *A* = 1*/*2, the system is in an intermediate regime between draft and drift — neither purely driven by selective sweeps nor by random sampling — which we call the drAIft regime (see Appendix A.4 for details). This contrasts with the loose draft regime (*A <* 1*/*2), where the effective population size is *N* ^2*A*^ ≪ *N*, reflecting a genuine change of time scale driven by selective sweeps, and with the drift regime (*A >* 1*/*2), where the coalescence probability is dominated by random sampling. We note that previous studies report a coalescence probability scaling as 1*/N* ^2*A*^ regardless of the value of *A*. Our finite size correction shows that for *A >* 1*/*2, this scaling breaks down and the coalescence probability remains of order 1*/N* (see Eq. (A.24)), consistent with a drift-dominated regime. Finally, we note that our second order approximation of Appendix A.4 breaks down at *A* = 1 (as in Barton (1998)) and more work would be needed to have a precise understanding of the drAIft regime for *A* between .5 and 1.

### 3.3 Loose genetic draft

#### 3.3.1 Effective population size

We now consider a population subject to recurrent selective sweeps. We assume, for the sake of simplicity, that each new beneficial allele appears at the same selected locus and benefits from the same coefficient of selection *s* over the resident allele. We shall further assume that successful selective sweeps occur at rate S. Following (Kaplan *et al*., 1989), one could consider two different cases, depending whether the adaptation process is mutation limited or not.

In the first case (mutation limited), the waiting time to the next successful selective sweep depends on the product of the number of new beneficial alleles per generation (*Nµ*, assuming a rate of beneficial *µ*) times its probability of fixation (well approximated by 2*s*, when *s* ≪ 1). This results in a rate of selective sweep of *S*_1_ ≈ 2*Nµs* that is proportional to *N* . For any fixed value for *s* and *µ*, the rate of selective sweep will increase to ∞ with *N* and the expected time between two selective sweeps will converge to 0. We do not consider this case here, where selective sweeps would, at the limit, occurred all nested one into the other and interfere. This would also result in a regime where *N*_*e*_ decreases as *N* increases as described in Weissman AND Barton (2012).

Alternatively, we consider here another population model where the rate of selective sweep does not depend on the population size and where an external factor allows for a new beneficial mutation to arise. An example of such regime is evolving in a fitness landscape with a moving optimum. Here we model environment changes at rate *λ* and the fitness optimum is shifted by *s*. After each change of environment, the population adapts to the environmental shift by a selective sweep.

If the time of adaptation is small compared to the time scale of the environmental changes, *λ* ≪ 2*Nµs* changes in the environment are almost instantaneously followed by a rapidly occurring selective sweep. In this setting, the sweep rate is then given by *λ* and is independent of *N*, a case that will always occur for sufficiently large values of *N* . In this second scenario, we have

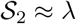

In the following, we will only focus on the case *S* ≈ *λ*. We will also assume that sweeps are not overlapping, which requires that

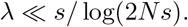

We monitor the fate of a neutral locus located at a map distance *c* of the selected locus. If we sample 2 individuals in the extant population under changing environment, we expect their coalescence to be driven by recurrent non-overlapping selective sweeps. We have shown in the previous section that the probability of coalescence between two lineages in a single sweep is well approximated (for *N* → ∞) by 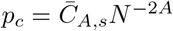, where

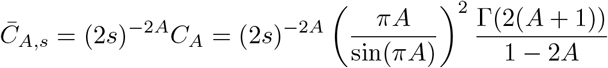

in the diffusion approximation (eq. A.10). It follows that the number of selective sweeps required for the two lineages to coalesce follows a geometric distribution with parameter *p*_*c*_.

Consequently, when *A <* 0.5, the expected time of coalescence between the two lineages at the neutral locus will be given by:

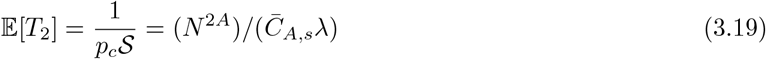

neglecting the effect of genetic drift (that occurs at a rate 1*/N*) that can be ignored for *A <* 0.5 (when *N* ^2*A*^ ≪ *N*).

A commonly used estimator of the effective population size based on the number of pairwise difference (Tajima, 1983), is proportional to the average 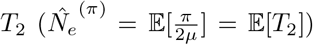. Using 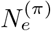, we come to the conclusion that for a locus subject to genetic draft with *A <* 0.5, the effective population size will no longer be linear in census size but a power function of it:

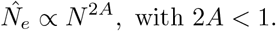

This implies that the time scale of the coalescent process is *N* ^2*A*^, which potentially will be much smaller that the census size *N* . The smaller the *A*, the stronger the reduction in coalescent time. Coalescent time remains nonetheless geometrically distributed, provided that *N* ^2*A*^ ≪ *N*, which will happen when *A* is not very close to 0.5.

To check the validity of this result, we have compared results from Monte-Carlo simulations (forward WF and backward RIF) to the predictions of equation (3.19), for the case of changing environments.

Forward simulations corresponds to a haploid Wright-Fisher model with a genome of two loci (one neutral one selected) for which the environment changes at an exponential rate *λ*. Beneficial mutations can occur only once per individual at the selected locus after an environmental change. The population evolves for a large number of generations (about 10*N* generations) where all parent-offspring relationship are recorded. At the end, two lineages are sampled at the onset of the last environmental change. Their ancestries are tracked backward until they coalesce.

Backward simulations corresponds to a continuous time backward RIF coalescent model with no overlapping selective sweeps. We follow two lineages sampled just before an environmental change. Environment change occurs at an exponential rate *λ* and the waiting time to the first selective sweep (after the change) is exponential with rate 2*Nµs*. If the later happens by chance to be longer than the former, an extra environmental change is considered. Lineages can coalesce during a selective sweep when they have the same type (as we did for simulations in Figure 3). Time is discretized in units of 0.001*s* during the sweep. Alternatively, lineages can coalescence between selective sweeps at a rate 1*/N* .

Results (figure 5) again show that forward and backward simulations matches closely for all values of *A*. For small *A* (in the figure *A* = 0.01), the effective population size is almost independent of *A*, almost 1*/λ*. For *A <* 0.5, the prediction of the simple approximation is good for large enough values of *N* . For smaller *N* values, the standard genetic drift dominates and simulations matches *N*_*e*_ = *N* . For values larger than *A >* 0.5, although we have no analytical approximation, the coalescent rate is on the order of *N*_*e*_ = *O*(*N*).

**Figure 5.**
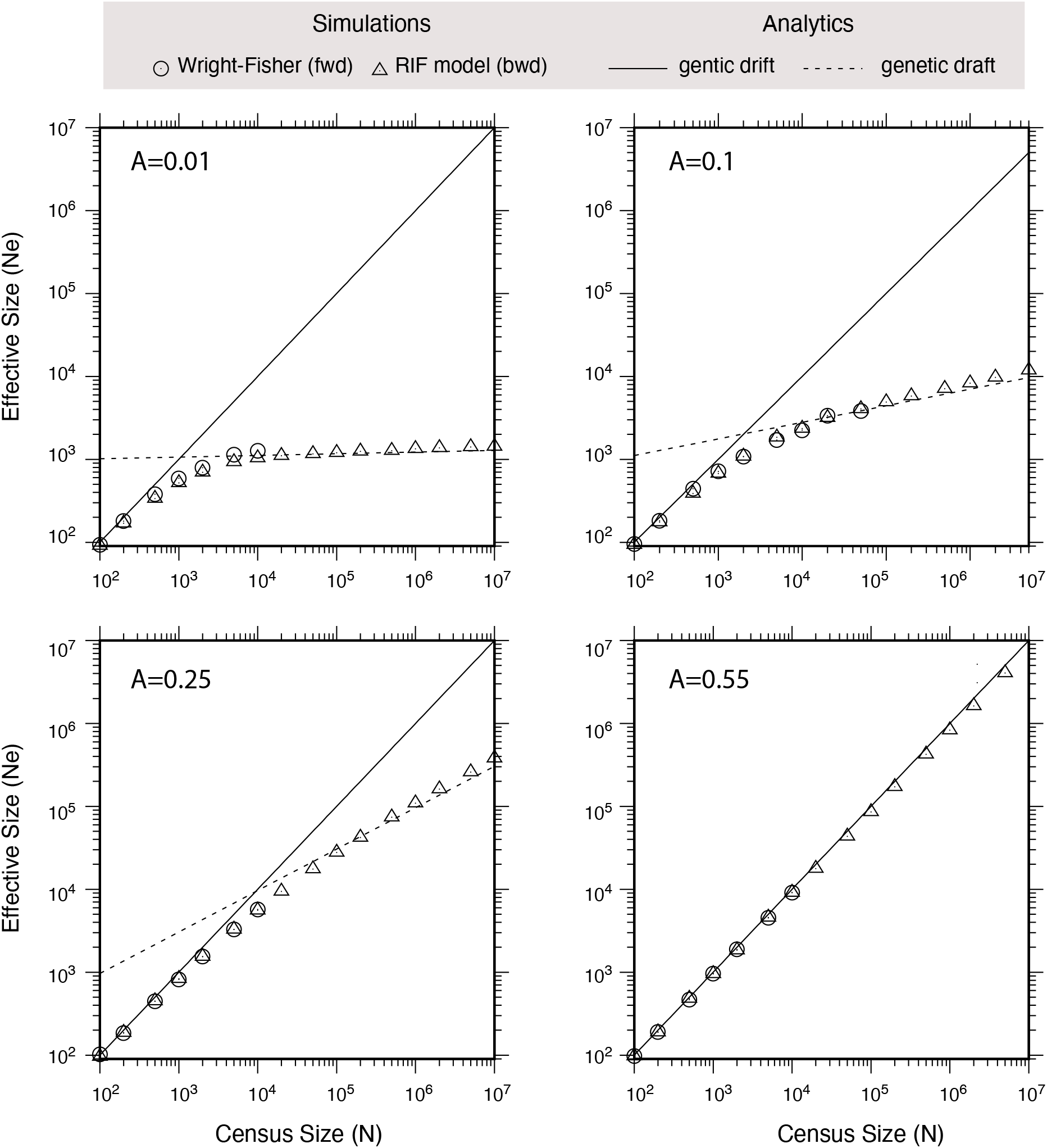
Average *T*_2_ is a power function of 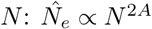, for *A <* 0.5. The solid line represents the neutral case *N*_*e*_ = *N*, and the dashed line represents analytical predictions (eq. 3.19). The circles are averages of 10^3^ forward individual based Wright-Fisher simulations. The triangles are averages of 10^4^ backward RIF simulations. A line in this log-log plot signifies a power law. Each panel is a different value for *A* = {0.01, 0.1, 0.25, 0.55}. The other parameters are *λ* = 0.001, *Nµ* = 0.1 and *s* = 0.05. On average, a sweep lasts 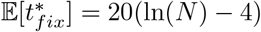 generations while the environment changes every 1*/λ* = 1, 000 generations.

We do note that the averages of *T*_2_ in the simulations are often slightly larger than the ones expected from equation (3.19) for *A* = {0.01, 0.1, 0.25}. This is mainly due to the existence in the simulations of few rapid environment changes where selective sweeps have no time to occur. In the theoretical predictions, any change of environment leads to a new selective sweep. Hence the small difference.

#### 3.3.2 Convergence to the Kingman tree

When *A* is large enough (for *A >* 0.1 in most of our simulations), the coalescence probability of two lineages for each selective event is small. In fact, it gets smaller as *A* gets closer to 0.5. As most of the selective events will not result in coalescing the two lineages, the fate of the neutral locus becomes mostly insensitive to the sweeps occurring at the selected locus.

Under loose genetic draft, we hypothesize that the probability of observing *k* lineages coalescing in a single selective sweep is much smaller than the pairwise lineages coalescence rate. We can thus define an *effective* population size

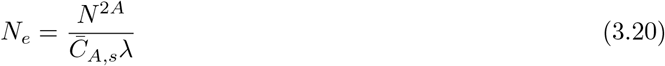

as the inverse of the expected pairwise coalescent time (equation 3.19). From there, the negligible probability of the multiple mergers coalescence events leads to a genealogy of *k*-sample that converges to a *k*-Kingman coalescent, once time is accelerated by *N*_*e*_. In particular, *T*_*k*_ the first coalescence time among *k* lineages is then approximately distributed exponentially with rate *k*(*k* − 1)*/*2*N*_*e*_.

To assess the convergence of the loose draft selective regime to a Kingman coalescent, we numerically compute the mean 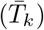 and the variance 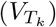 of *T*_*k*_ for *k* = 2, · · ·, 7. If the tree is a *k*-Kingman, 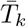 and 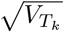 should both converge to 2*N*_*e*_*/k*(*k* − 1). As a visual test, we compare 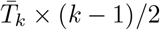 and 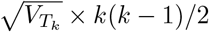 to see whether it converges to a constant (that should be *N*_*e*_).

Results (Figure 6) shows that means and standard deviations of *T*_*k*_ for *k* ∈ [2, 7] are usually close numerically supporting the convergence to exponential distributions. Furthermore, they are usually also in fairly good agreement with our theoretical expectations (genetic drift for small sizes, e.g. *N* = 100 and genetic draft for large population sizes *N >* 10^5^), visually supporting the convergence to an approximately *k*-Kingman tree.

**Figure 6.**
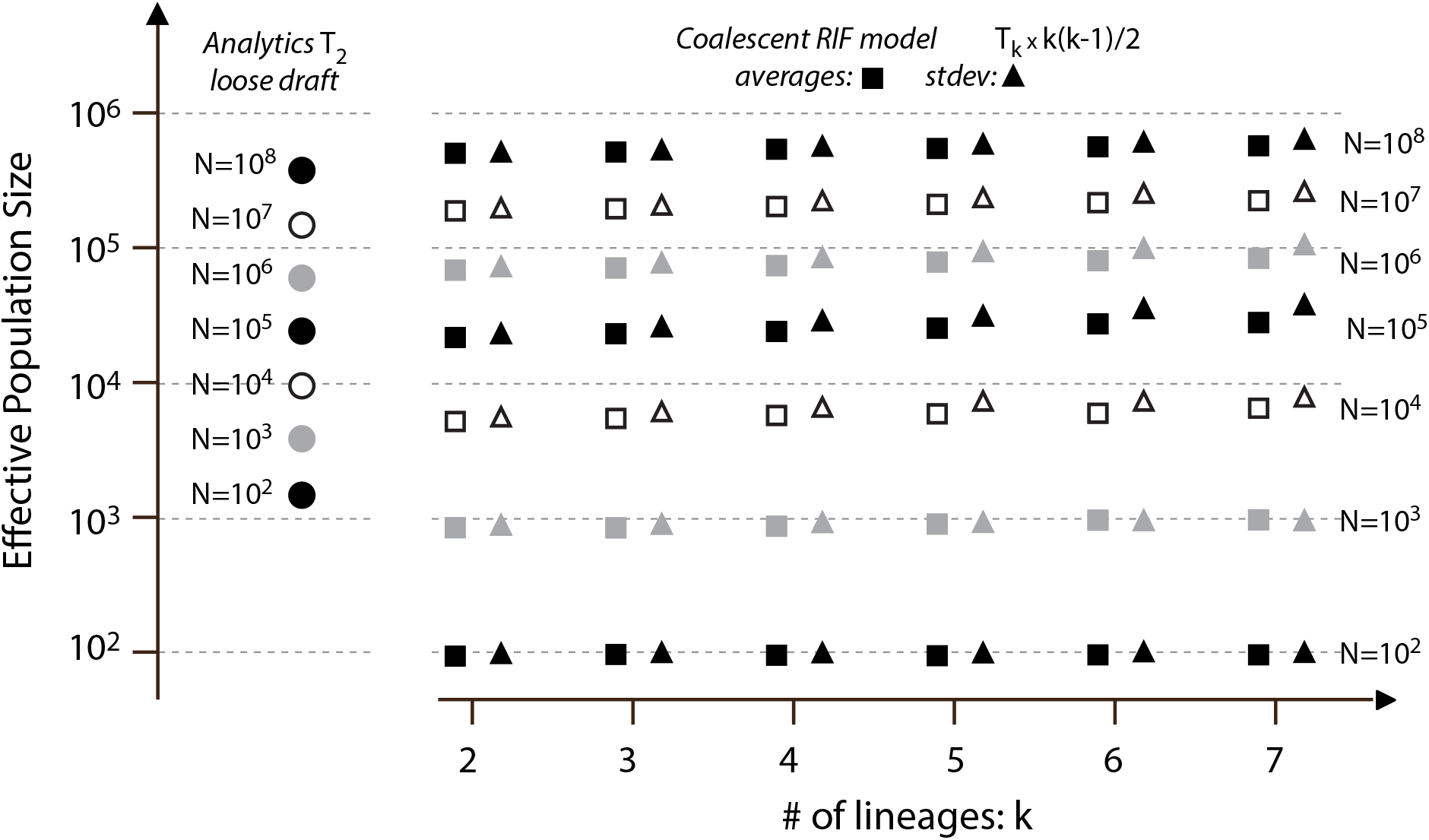
Time scale (*N*_*e*_) of each stage of the coalescent tree. For large sizes, *N >* 10^4^, the averages and their associated standard deviations follow approximately a Kingman coalescent distribution, but with a highly reduced *N*_*e*_. For all values of *N*, we have *c* = 0.003 and *s* = 0.01, so that *A* = 0.3.

This implies that under loose draft regime, the coalescent tree likely converges to a neutral Kingman tree, with an overall reduced time scale.

#### 3.3.3 Convergence to the Kingman SFS

An important feature of the neutral model is the Site Frequency Spectrum (SFS) that is the distribution of frequencies among the variants. Assuming a an infinite site model, the expected SFS is given by *E*[*ξ*_*i*_] = 2*Nµ/i* (Fu, 1995). Using the Monte-Carlo backward RIF simulator, we computed the average sample SFS for several values *A* for a population of *N* = 10^6^ experiencing recurrent selective sweeps at rate *λ* = 10^−4^. The SFS are reported first (top row, *ξ*_*i*_) in branch length of coalescent time scale (*N* generations), disregarding the mutations themselves and then (bottom row,, *φ*_*i*_ = *i* × *ξ*_*i*_*/* ∑*ξ*_*i*_) after normalization and transformation (Achaz, 2009) that results in a flat horizontal line for the case of Kingman coalescent.

Results (Figure 7) clearly shows that for *A* = 0, the expected SFS is monotonically rapidly decreasing, with almost exclusively singletons. If one assumes that no coalescence occur before the first selective sweep, one expects 20 × 10^4^ generations in external branches, resulting in *E*[*ξ*_1_] = 0.2 for the singletons. At the other extreme, in the Kingman regime (*A* → ∞), one expects *E*[*ξ*_1_] = 2.0 for the singletons (Fu, 1995).

**Figure 7.**
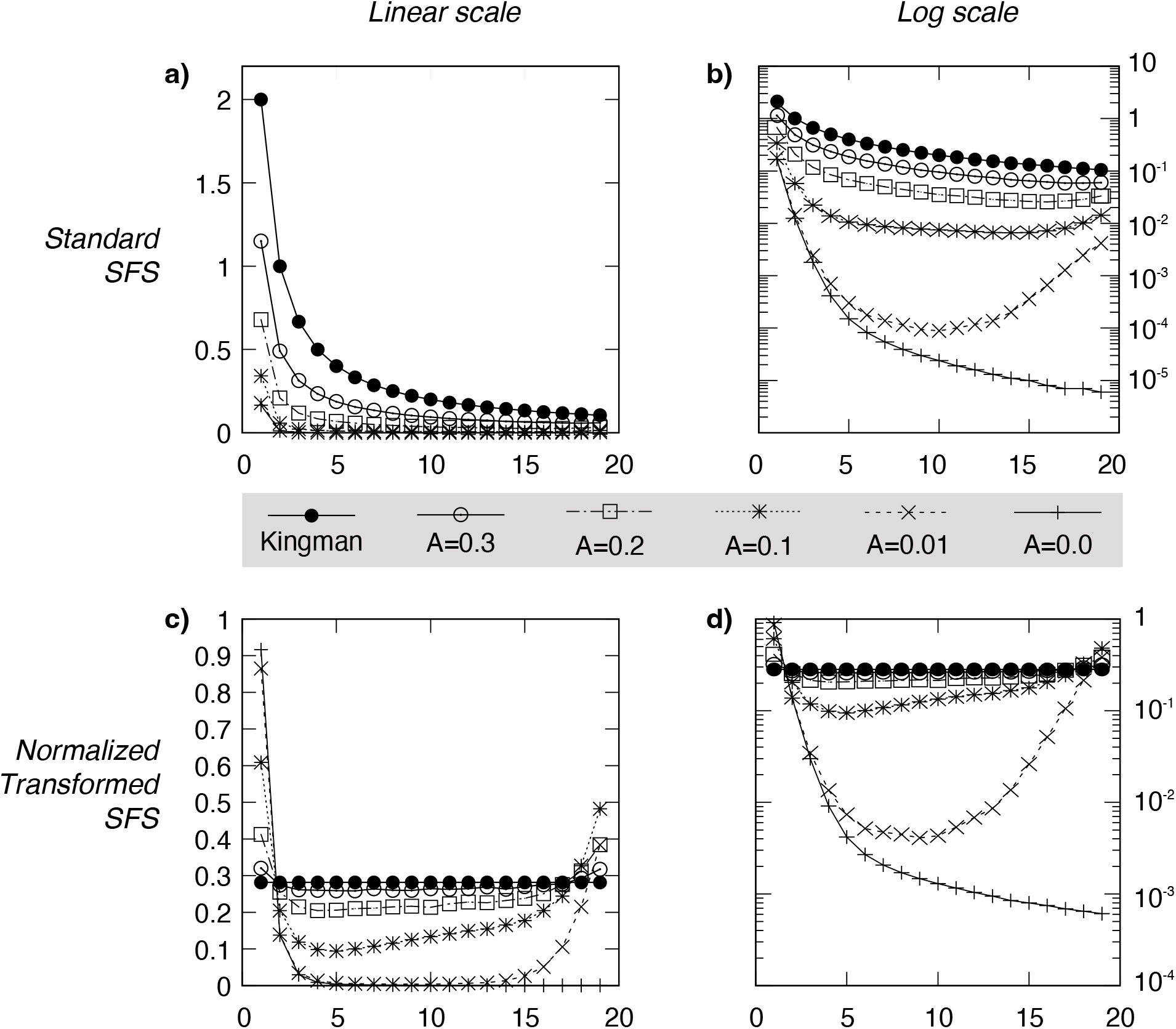
Average Site Frequency Spectrum (SFS) for various distance to the selected site. Averages were obtained using the backward coalescent simulator, with *N* = 10^6^, *s* = 0.01, *n* = 20, *λ* = 10^−4^, 10^5^ replicates and *A* ∈ {0, 0.01, 0.1, 0.2, 0.3}. The case of Kingman (*A* → ∞) is computed analytically. In the top row (a and b), the standard SFS (*ξ*_*i*_) is computed in units of branch length (in a coalescent time scale of *N* generations). In the bottom row (c and d), the SFS (*φ*_*i*_) is normalized and then transformed: *φ*_*i*_ = *i* × *ξ*_*i*_*/* ∑*ξ*_*i*_, resulting in an horizontal line for a Kingman coalescent.

Both draft regimes are well illustrated by the height and shape of the SFS. In the tight draft regime (0 *< A <* 0.1), the SFS exhibits a strong depletion of variants together with a strong U-shaped SFS (see the case of *A* = 0.01). For the loose draft regime (0.1 *< A <* 0.5), the SFS exhibits a clear depletion of variants but the shape of the SFS is Kingman-like (see *A* = 0.3). In this latter case, a neutrality test based on the SFS such as Tajima’s D (Tajima, 1989) or any other SFS-based neutrality test (Achaz, 2009) will hardly reject neutrality.

## 4 Discussion

In this article, we have studied the effect of recurrent positive selection on neutral loci located at a given map distance *A*, expressed in Morgans per unit of *s*. We have compared theoretical derivations with individual based forward simulations and with backward coalescent simulations. We have shown that the trajectory of the frequency of a selected allele for a single selective event in finite population is very well approximated by the RIF model (Martin and Lambert, 2015), a semi-deterministic proxy of a stochastic selective sweep where the initial and final stochastic phases are simplified to two instantaneous stochastic jumps. Beyond its accuracy, this approximation proved to be a key methodological asset: it is approximately 10^3^ times faster than a naive Wright-Fisher simulation, making it tractable to explore the large parameter space of loose genetic draft that would otherwise be prohibitively slow.

For the RIF model, we have derived an approximation for the coalescent probability under a strong selective sweep (*Ns* ≫ 1). Even the RIF model may appear as an over-simplistic representation of a stochastic sweep, we found that the latter approximation is consistent with a more complex diffusion model, and with a former corrected approach of BARTON (1998). It is also notably better than many previously derived approximations for “large” values of *A* (e.g. *A* ∈ [0.1, 0.45]) (see Figure 3). Based on this analytical approximation, we characterized the expected coalescent rate in a regime of recurrent sweeps (genetic draft regime) that result in reduced diversity *N*_*e*_ = *O*(*N* ^2*A*^) ≪ *N*, where *A* is the recombination/selection ratio. Importantly, under the draft regime, the relationship between *N*_*e*_ and *N* is not anymore linear, pointing the importance of deriving models with no proportionality between genetic diversity and population size *N* . As the effect of a single selective event is loose for large enough values of *A*, we named this regime loose genetic draft. This regime of loose genetic draft produces patterns that mimic neutrality and thus can be therefore framed in terms of *emergent neutrality* (Schiffels *et al*., 2011; Neher and Shraiman, 2011). Importantly, at the limit, the mimicking is perfect in the loose draft as it generates an n-Kingman tree, where nodes are entirely driven by the selective events.

Our predictions differ somehow from previous models of linked selection that have focused on different types of regime for the effect of linked selection. A number of studies (Etheridge *et al*., 2006; Durrett and Schweinsberg, 2004; Weissman and Barton, 2012; Coop and Ralph, 2012; ELYASHIV *et al*., 2016) have focused on the impact of tight genetic draft, for which the distance to the selected locus is small (*A* ≪ 1). Let us now emphasize one important difference between tight and loose genetic drafts. In the tight regime (*A* ≪ 1), selection is so large as compared to recombination that the coalescence probability at every sweep is non-negligible. As a consequence, the time to coalescence is dictated by the time scale of the time elapsed between two consecutive sweeps.

Following up on our earlier discussion on recursive sweeps, we can explore a scenario where sweeps are triggered by environmental changes followed by rapid adaptations. As before, if we denote by *λ* the rate of environmental change, the time to coalescence is in tight draft typically of order *O*(1*/λ*), *regardless of the population size*. Thus, in a tight genetic draft scenario, theory predicts that genetic draft could potentially lead to a reduction of *N*_*e*_, but this reduction is problematic since it completely decorrelates *N*_*e*_ from *N* . This phenomenon is illustrated on the first panel of Figure 5 where *N*_*e*_ is almost insensitive to *N* . This important limitation of the tight draft hypothesis was already raised in previous works (Coop and Ralph, 2012; Buffalo, 2021; Charlesworth and Jensen, 2022) and has been advocated as one argument against the hitchhiking effect as a potential explanation for the observed discrepancy between 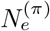 and *N* .

One contribution of our work is to show that a loose hitchhiking effect (loose draft) overcomes this limitation and generates a power law relation between *N* and *N*_*e*_. For 0.1 *< A <* 0.5, this is illustrated on Figure 5 for both simulations and our analytical predictions. This dependence arises from the dependence in *N* of the pairwise coalescence probability during a single sweep. In the tight draft regime (*A* ≪ 1), this probability is barely sensitive to *N* . As the coalescence probability is 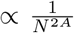 so that when *A* = *O*(1) the effective population size is 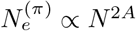.

We would like to emphasize that this seemingly technical point has lead to some confusion in the past. The loose draft regime has been alluded in an influential article by COOP AND RALPH (2012) where it is stated that the expected pairwise difference satisfies (see equation (18) therein)

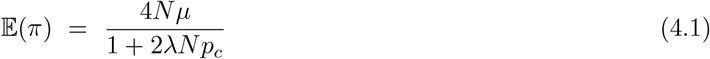

where *µ* is the mutation rate at the neutral locus and *p*_*c*_ is a parameter which encapsulates the pairwise coalescence probability during a single sweep. Based on (4.1), Coop and Ralph concludes that “the key parameter is the population-scaled rate of sweep 2*λN* “. In particular, this lead to the conclusion (see e.g. (Buffalo, 2021; Charlesworth and Jensen, 2022)) that the loose recursive selective sweep would generate an effective population size *N*_*e*_ independent of *N* as along as 2*Nλ* ≫ 1. An important difference between the approach in (Coop and Ralph, 2012) and the present work lies in the assumptions on the coalescence probability *p*_*c*_. In (Coop and Ralph, 2012), *p*_*c*_ is considered as an exogenous parameter reflecting some hidden sweep mechanism (e.g., random selection coefficient, changing environment during the sweep). This is in contrast with the present work where we start from an explicit individual based model and we show that the coalescence probability *p*_*c*_ has an implicit dependence in *N*, i.e. 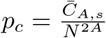. Plugging this expression into (4.1), we find now that for *A <* 0.5 (neglecting the first term of genetic drift in the numerator):

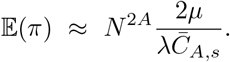

Again, this leads to an estimation of the effective population 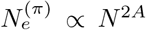 which is consistent with our previous estimation of E(*T*_2_) in eq. (3.19). To summarize, our approach is consistent with the one of Coop and Ralph, but we provide an explicit coalescence probability as a function of the population size *N* . In turn, this leads to a drastically different estimation of the effective population size.

In addition to the effect on the coalescence time scale, different regimes of genetic drafts result in different types of genealogical structure. As already discussed, the limiting genealogy in a loose draft regime is indistinguishable from a neutral genealogy (Kingman coalescent). In contrast, for tight draft scenario, the linkage is so tight that typically, several lineages coalesce at each selective sweep, resulting in processes that are known as Multiple Merger Coalescents (MMC) (Berestycki, 2009; Tellier and Lemaire, 2014) . At the limit of *A* = 0, the genealogical tree of the neutral is entirely driven by the last selective event. If this event was strong *Ns* ≫ 1, the genealogy reassembles a radiation where all lineages coalesce very rapidly. For slightly larger values of *A*, only a part of the lineages coalesce at each selective events resulting in trees such as Bolthausen-Snitzman trees or more generally other MMC such as Λ − *coalescent* or Ξ − *coalescent*.

On figure 8, we have pictured the different regimes of evolution as a function of the distance to the selected loci.

**Figure 8.**
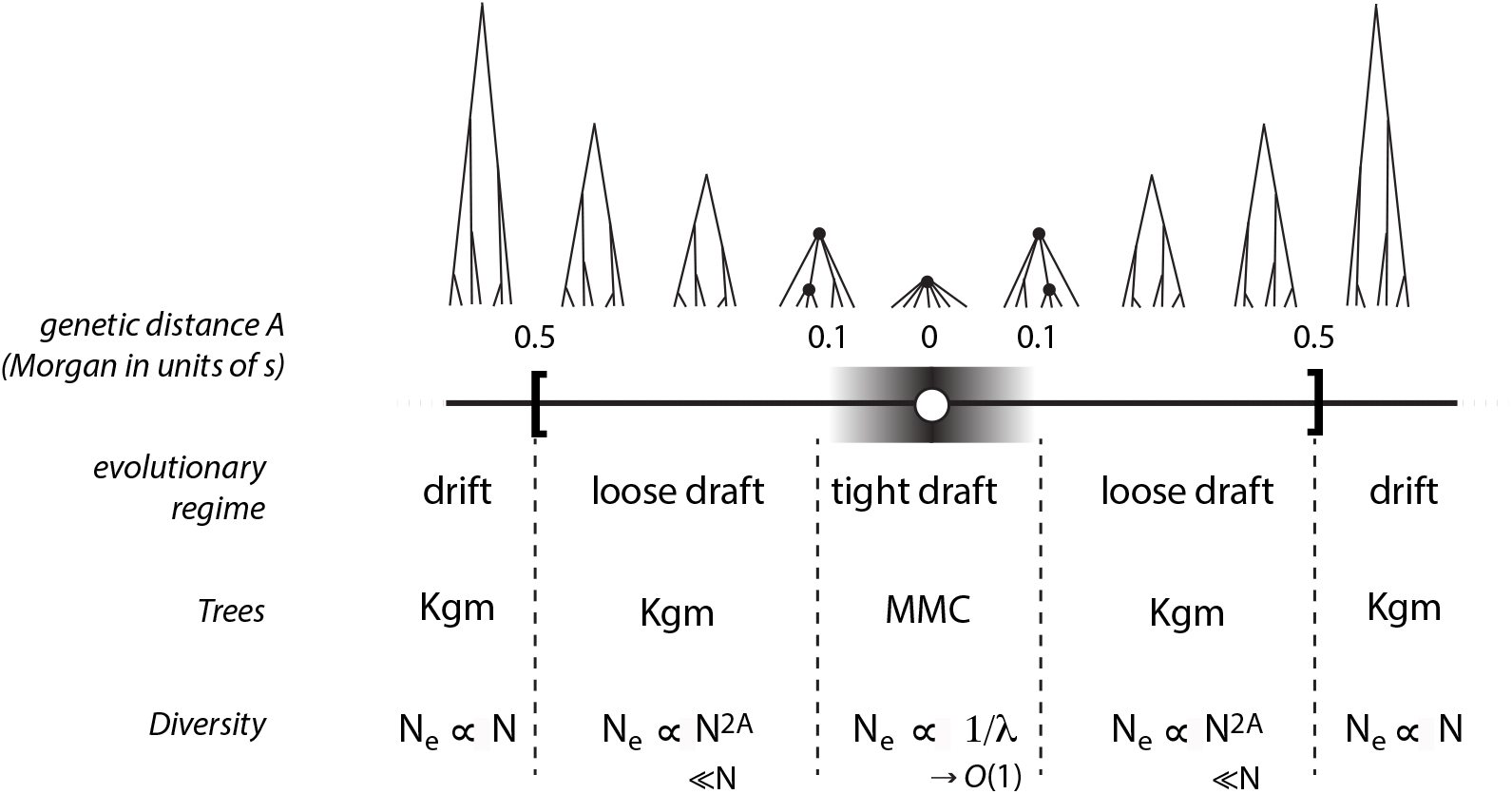
Overview of the different regimes of evolution as a function of the distance to the selected locus. **tight draft**: at small distance (*A* ≈ 0), the tree is best described by a Multiple Merger Coalescent and the diversity is insensitive to population size: 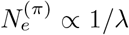. **loose draft**: at larger distance (*A* ∈ [0.1, 0.5[), the tree is best described by a Kingman tree but diversity is highly reduced 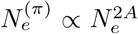. **Genetic drift**: at even larger distance (*A* ≥ 0.5), the tree is a Kingman tree and diversity is given by the population size: 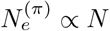.

- At small distance (*A <* 0.1), the neutral loci are highly impacted by the selected events (tight draft regime) : (a) their genealogy is described by a Multiple-Merger Coalescent (MMC) tree (Berestycki, 2009; Tellier and Lemaire, 2014) and (b) their genetic diversity is extremely reduced and *N*_*e*_ is insensitive to the underlying population size *N* (*N*_*e*_ ∝ 1*/λ*).
- At intermediate distance (0.1 *< A <* 0.5), the neutral loci are moderately impacted by the selected events (loose draft regime) : (a) their genealogy is described by a Kingman (Kgm) tree although (b) their genetic diversity is largely reduced 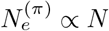.
- At large distance (*A >* 0.5), neutral loci are weakly impacted by selective events (genetic drift regime): (a) their genealogy is described by a Kingman (Kgm) tree and (b) diversity *N* ^(*π*)^ ∝ *N* .
- Finally, close to *A* = 1*/*2, we identified an intermediate regime (the drAIft regime) where the genealogy is Kingman and where the effective population size is only reduced by a constant relative to *N* .

Intuitively, if one assumes that beneficial mutations arise frequently and are scattered throughout the genome, the average genomic pattern will likely be the result of a mix of the different scenarios, where the strongly affected regions likely encompass only a small proportion of the genomes. It is then tempting to postulate that the average patterns of diversity such as genome-wide SFS would be better fitted by Multiple Merger Coalescent models than by a pure Kingman coalescent model, an observation recently supported for a wide range of organisms (FREUND *et al*., 2023; ÁRNASON *et al*., 2023).

Two different regimes of genetic draft can be envision. In a first scenario, each new selective sweep occurs at a uniformly randomly chosen site in the chromosome. In this regime, the strongest diversity reduction for a neutral site, will be driven by rare events where the selected sites is closely linked to the studied neutral locus. The loose draft regime in this scenario only occurs transiently until a beneficial variant appears in the vicinity of the neutral locus. In the second case, potentially selected sites have fixed positions and thus neutral locus can well be always at a appreciable map distance of the selected site. For these neutral loci, the regime of loose genetic draft will entirely drive their evolution.

Regardless of the exact draft regime, neutral loci affected by selection should be in the segment of size *A* (0.5 on both sides) of the selected loci. To have a crude sense of theses distances, if we consider selected events of *s* = 0.01 (1.01 relative advantage), the impacted area of a selected events is 1 cM (about 1Mb of the human genome and more generally on the order of 1% of a chromosome). Stronger selected events will impact the chromosome on a larger region. At any generation, many selective events can occur without driving the species to extinction (hundreds for Haldane (1957), much more for Maynard Smith (1968)). Therefore, it is likely a large fraction of the genome is affected by linked selection at any generation, an hypothesis that has been proposed based on different arguments (Pouyet *et al*., 2018).

One restrictive assumption of the current study is that selective sweeps are non-overlapping. For the loose draft regime to have a sizeable effect on the effective population size, the sweep rate *λ* must satisfy

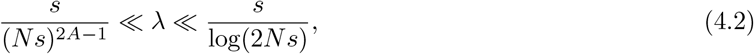

which requires *Ns* to be sufficiently large for this window to be wide enough in practice. We conjecture that the second inequality could be relaxed to allow for overlapping sweeps. It is therefore of fundamental importance to understand the loose draft regime in a scope of parameters where several beneficial mutations are in clonal interference. Whether our conclusions remain consistent with the present study (emerging neutrality together with a highly reduced time scale) and whether loose sweeps at higher frequency could potentially impact a larger scope of the genome (*i*.*e*. a recombination-selection ratio *A* beyond 0.5) requires to go beyond the current theoretical framework and is the subject of a further study.

Although the model discussed here is framed in terms of fluctuating environment that triggers selective sweeps, it is also compatible with a regime of evolution by compensation, where deleterious substitutions accumulate in the population until a compensating mutation appears and then sweeps to restore the original fitness (Kimura, 1985). The role of compensatory mutations in molecular evolution has since then been extensively supported by data analysis (e.g. (Kondrashov *et al*., 2002; Samecz *et al*., 2014; Latrille *et al*., 2024)). Compensatory evolution differs from the regime of background selection (Charlesworth *et al*., 1993; Matheson and Masel, 2024) by the recurrent triggering of new selective sweeps.

Effective population size estimates derived from diversity data are typically far below what the Standard Neutral Model would predict 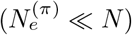. Nevertheless, a handful of life history traits have been identified as meaningful predictors of diversity. Species producing small propagules, whose abundance likely scales inversely with population size, tend to exhibit higher levels of diversity (Romiguier *et al*., 2014). More recently, analysis of a 172-species dataset provided strong statistical evidence for a positive log-log linear relationship—that is, a power law—between census size and genetic diversity (Buffalo, 2021).

Because neutral diversity scales with coalescence time, captured by 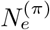, Lewontin’s paradox can be reframed as a puzzle about the timescale of genetic ancestry. Resolving it therefore requires explaining the order-of-magnitude gap between 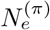 and *N* . Non-constant demography, population structure, and linked selection can each drive 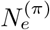 far from *N* . However, most departures from the Standard Neutral Model proposed to account for reduced diversity also produce pronounced distortions in other diversity statistics, such as the site frequency spectrum (SFS). Moreover, many of these mechanisms predict an effective population size that either remains on the order of *N* (e.g., age structure or skewed sex ratios; WRIGHT 1931), or becomes nearly independent of *N* altogether (e.g., tight linked selection or severe demographic fluctuations; Coop and Ralph 2012).

In this study, we demonstrate that loose draft can yield an effective population size that scales as a power law of the census population size, consistent with empirical findings (Buffalo, 2021). Crucially, unlike many previous mechanisms, loose draft does not distort standard diversity statistics such as the SFS. In brief, under the loose draft framework, neutral sites retain quasi-neutral diversity patterns while the effective population size scales as a power of the census population size.

Although, we do not pretend that this simple loose genetic draft model is realistic enough to solve Lewontin’s paradox of variation, it nonetheless is an interesting step forward in understanding the effect of sparsed linked selection and in proposing models where diversity does not scale linearly with population size but as a power law that linear in log-log, an observation strongly supported by the most recent data analysis (Buffalo, 2021). Although the model we have consider here is simplistic as it assumes no demography, no structure and fixed values of *c* and *s*, it paves the way for future more realistic models. It now remains to explore all relaxations of the model such as having overlapping selective sweeps, random *c* and *s*, soft selective sweeps, incomplete sweeps to appreciate how it tunes diversity as a function of population size.

## Acknowledgments

We would like to thank F. Ged and N. Barton for providing constructive feedbacks on this manuscript. We also thank D. Weissman for suggesting the general solution to the ODE in BARTON (1998) and its relation to our diffusion approximation.

## A Appendices

### A.1 Coalescence probability under the RIF model

Consider now two lineages starting in state 0. Let *T*_*c*_ be the time to coalescence which is possibly infinite if the two lineages do not coalesce. Our aim is to compute:

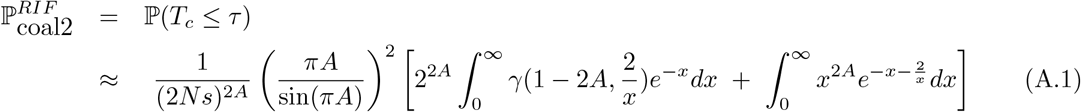

The derivation of this formula is decomposed below in several steps.

#### A single lineage

We first consider a single lineage starting in state 0 at time 0. We first show that the probability of a single lineage to be in state 0 close to time *τ* (time *τ* − *t* close to the onset of the sweep forward in time) is of the order 1*/*(*Ns*)^*A*^. We recall that 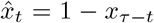 is the reversed RIF trajectory. We consider the conditional probability of being linked to a type 0, *u*_*t*_ := ℙ (*L*_*t*_ = 0|ℰ, ℰ′). Then

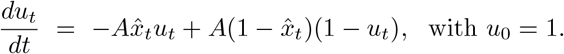

A direct computation around *τ* leads to

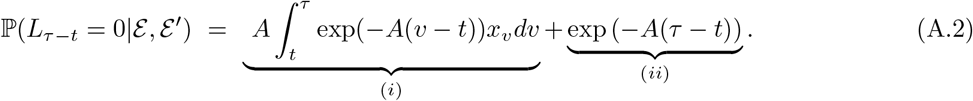

We first evaluate the term (i)

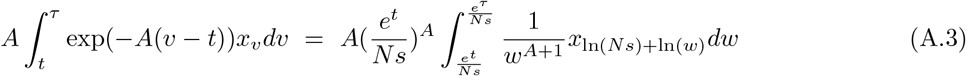

where we used the change of variable *w* = *e*^*v*^*/Ns*. From the explicit solution of the logistic equation for *x*, this yields the limit

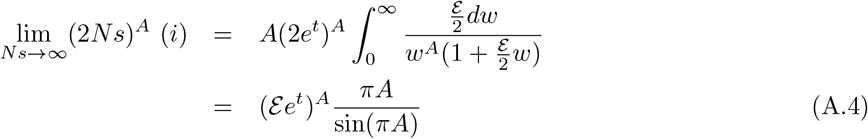

We now show that the second term (*ii*) is negligible compared to (*i*) as *Ns* ≫ 1. This is indeed the case since, as *τ* is given by (3.11), we have

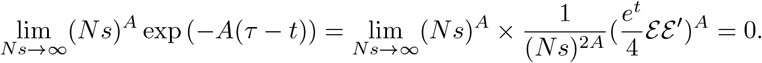

This implies that (*ii*) = *o*(*i*). Gathering the previous estimates yield

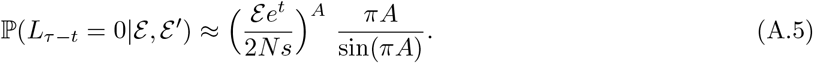

#### Two lineages

We will now use the previous estimate to evaluate the coalescence probability of two lineages during a sweep. Each lineage starts at backward time 0 with a type 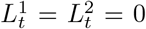 and can switch to state 1 by recombination at rate 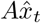. It switches back to type 0 at rate 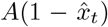. Two lineages of type 0 coalesce at rate 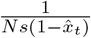 and they coalesce at rate 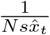 when both are type 1. Figure 9 provides an illustration of the notations of the considered process.

**Figure 9.**
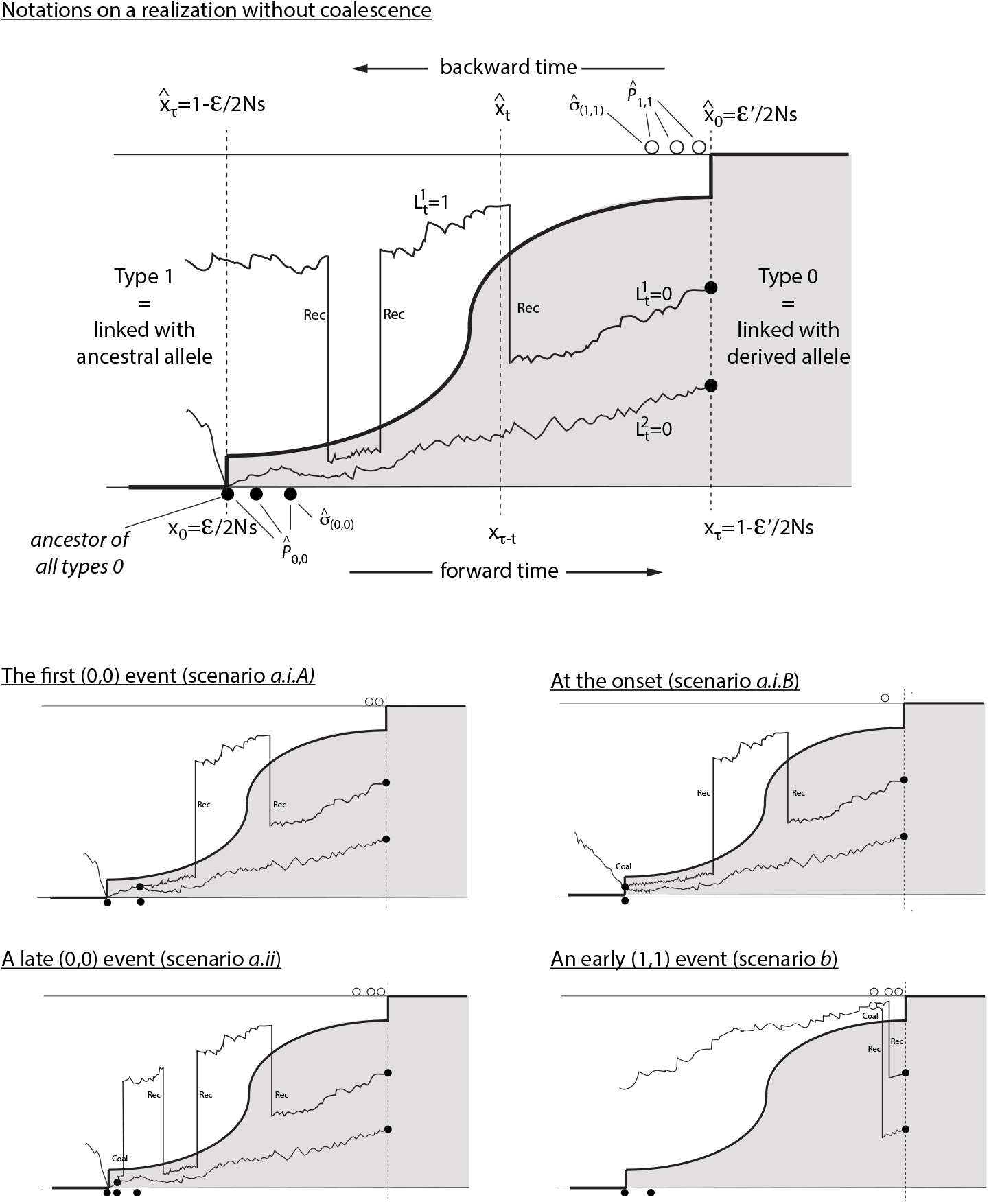
A sketch of the coalescent process under the RIF model. The top panel presents all notations used in the text. The four bottom panels reports all possible scenarios for an effective coalescence. The top panels represent the most likely events, namely when the two lineages are type 0 at the first (0, 0) potential coalescence event 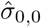. **scenario** *a*.*i*.*A*: the coalescence occurs at time 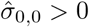. **scenario** *a*.*i*.*B*: no *positive* potential (0, 0)-coalescence times. Coalescence occurs at the onset of the sweep at backward time *τ* . (part B of (A.1)). **scenario** *a*.*ii*: There is a (0, 0)-effective event occurring after the first potential (0, 0)-event 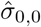. In this scenario, a lineage transitions from 1 to 0 on the time interval 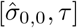. **scenario** *b* effective (1, 1) coalescence event. The two lineages must transition from 0 to 1 before the occurrence of the last (1, 1) potential coalescence event 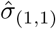.

We will now show that under the RIF model, the probability of coalescence during the selective sweep is well approximated by a sum of two terms:

i. the probability of coalescence between two types 0 at *t* = *τ* − *O*(1), close to the onset of the selective sweep.
ii. the probability that both lineages are type 0 and reach *τ* because they had no opportunity for coalescence before. They immediately coalesce in the ancestor of all derived alleles.

To get our approximation of the coalescence probability, we construct the pair of *dependent* coalescing lineages 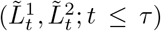 from two *independent* lineage processes 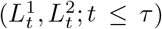. This is done as follows. Given a realisation of the reversed fixation trajectory 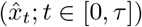, we define two independent Poisson Point Processes 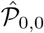 (potential coalescence events between two types 0) and 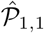 (potential coalescence events between two types 1) with respective intensity measures:

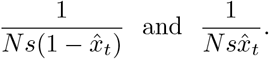

To enforce the constraint that (0, 0)-lineages coalesce at time *τ*, we add an extra point to 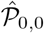 at time *τ* .

Next, we define *T*_*c*_ as the time at which a potential coalescent event 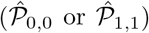 coincides with two lineages having the correct type 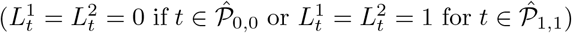. At such time, the two lineages must coalesce. We call such a potential event an effective coalescence event. Formally:

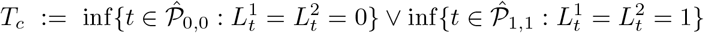

with the possibility that it never occurs. Finally, we define a pair of *dependent* coalescing lineages as identical to the *independent* ones before *T*_*c*_, while they both equate after *T*_*c*_. Formally:

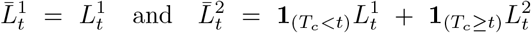

Using this construction, we will now argue that the coalescence probability of two lineages is approximately the probability that *L*^1^ and *L*^2^ are in state 0 at the first potential coalescence event 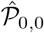. According to the previous construction, coalescence can be decomposed into 3 scenarii (see *a*.*i, a*.*ii* and *b*). Figure 9 provides an illustration of all possible coalescence events.

**scenario** *a*. The effective coalescence is a (0, 0)-event. This scenario is itself decomposed into two sub-scenarii (see below). First, let 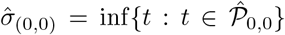 be the first potential (0, 0)-event. Regarding 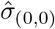, an effective (0, 0)-event can only occur in two ways:

**scenario** *a*.*i*. The independent lineages (*L*^1^, *L*^2^) are in state (0, 0) at time 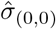. Either they coalesce at 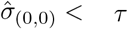 (scenario *a*.*i*.*A* of Figure 9), when the first (0,0) event is before the onset on the sweep, or at 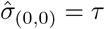 when no (0,0) event exist before the onset (scenario *a*.*i*.*B*).

**scenario** *a*.*ii*. The coalesce occurs at some time 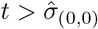, that is at subsequent point 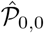, closer to *τ*, the onset of the sweep.

**scenario** *b*. The coalescence is a (1, 1)-event. Both lineages recombine with resident alleles very early and coalesce at one of the (1,1)-event.

We will now show that only scenario *a*.*i* contribute significantly to the probability of coalescence and that the scenarii *a*.*ii* and *b* can be neglected.

**Scenario** *a*.*i*: This probability can be decomposed into two parts:

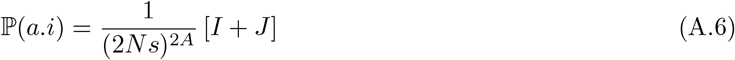

where

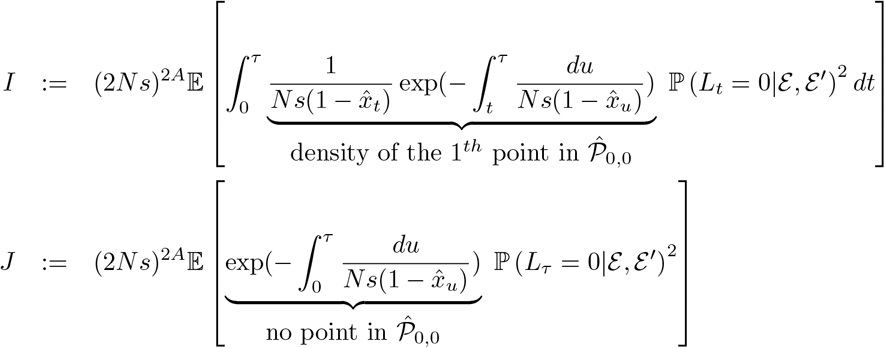

Here, *J* is the (rescaled by (2*Ns*)^*A*^) probability that the only potential (0, 0)-event is time *τ* (onset of the sweep forward in time); and *I* is the (rescaled) probability that there is at least one potential (0, 0) event different from *τ*, and that the independent lineages are in state 0 at the first potential (0, 0) event.

To estimate *I*, we use that for a fixed time *t*,

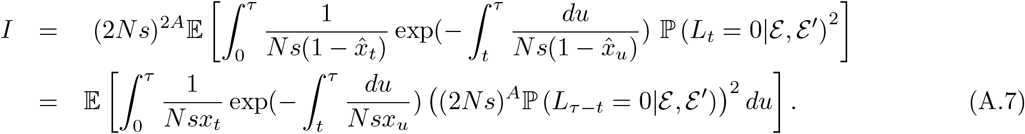

We have 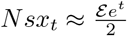 when *Ns* → ∞. Thus

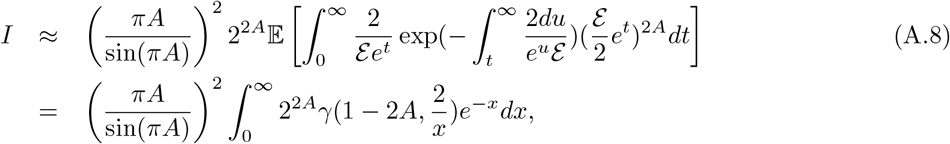

where *γ* is the lower gamma function. The second line is obtained by the change of variable 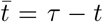. In the third line, we used the approximation 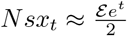. By a similar computation, we estimate *J* and further show that:

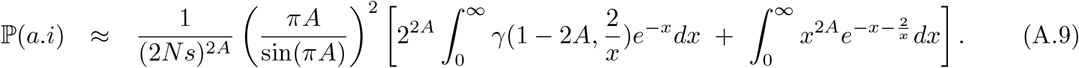

To derive our final result (A.1), we now show that scenarii (*a*.*ii*) and (*b*) are negligible compared to scenario (*a*.*i*). Let us first estimate the probability of scenario *b*, that is a (1, 1)-coalescence event. For a (1, 1)-effective event, at least one of the two independent lineages transitions from state 0 to 1 in the time interval 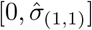, where 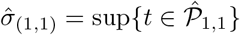. This entails that

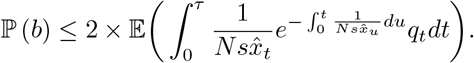

where *q*_*t*_ = ℙ_0_(∃*u* ∈ [0, *t*], *L*_*u*_ = 1|ℰ, ℰ′). Now,

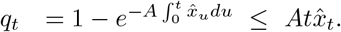

It follows that

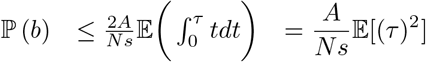

Since *τ* = *O*(ln(*Ns*)), it follows that

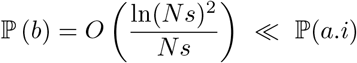

It remains to estimate scenario (a.ii) that coalescence occured at a (0,0) event after 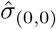. Under this event, one of the two lineages transitions from state 1 to 0 on the time interval 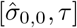. By reversibility of 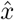, it is not difficult to see that the probability of latter event is identical in distribution to our upper bound for (b), so that

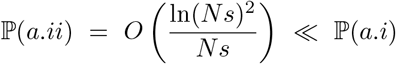

Gathering all the previous estimates, this yields the coalescence probability as written in (A.1).

### A.2 Coalescence probability under the Diffusion model

#### A first derivation

A similar reasoning on the diffusion model shows that the pairwise coalescent probability can be approximated in the following way

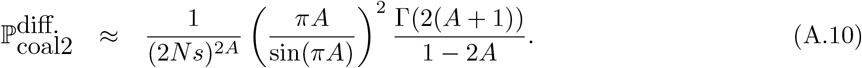

The rational behind this result is a combination our the approach developed in the RIF model together with the method developed in (Etheridge *et al*., 2006). We decompose the outline of the proof into 4 steps.

##### Step 1

As in the RIF model, we start from the observation that the probability of coalescence can be decomposed approximately as in (A.6) where the term *J* is now 0 since the process is continuous and starts at 0. For the term *I*, the approximation (A.7) derived for RIF is also valid in the diffusion case, that is, one can show along the same lines that

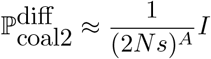

where

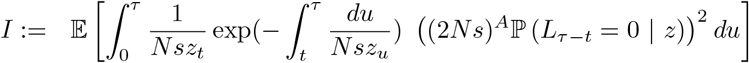

where *z* is solution of the Stochastic Differential Equation (3.4). As in RIF, this is justified by the observation that the probability of coalescence is well approximated by the probability of finding the two independent ancestral lineages in the 0 population at the first point of potential coalescence time in the point process 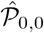. We will not repeat the arguments here and refer the reader to Figure 9 for reference.

##### Step 2

Recall that for a fixed time *t*, the process *Nsz*_*t*_ is well approximated by the process *y*_*t*_ as defined in (3.7). As in (3.8), we define ℰ as the stochastic correction to the exponential growth of the population process *y*

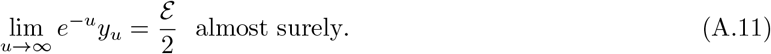

By arguing analogously to the RIF model (see the reasoning yielding (A.5)), we can approximate *Nsz*_*t*_ by *y*_*t*_ in the previous integral and

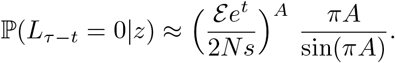

The arguments are completely analogous to the RIF case and will be skipped for the sake of conciseness.

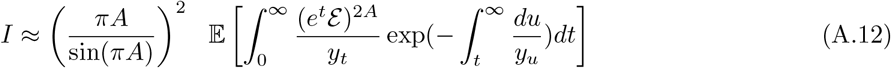

##### Step 3

We are now left with the previous expectation whose integrand only depends on the population process *y*. Recall that *y* can be thought of as the continuum limit of a near-critical branching process conditioned on non-extinction. In order to compute the latter quantity, we will rely on the genealogical structure of the diffusion *y* that we now describe. To make this genealogical structure explicit, we start from a Yule process *Y* starting from a single individual with branching rate 1. We think of *Y* as describing the infinite lines of descent in the conditioned supercritical continuum branching process *y*. In (Evans and o’connell, 1994), it is demonstrated that the process *y* can be constructed by allowing “mass” to emigrate from the ancestral lineages. A visual illustration of how the process *y* can be reconstructed from the ancestral process *Y* is provided in Fig. 10. Intuitively, the mass that departs from the genealogy *Y* can be interpreted as the descendants of a particular infinite ancestral line.

**Figure 10.**
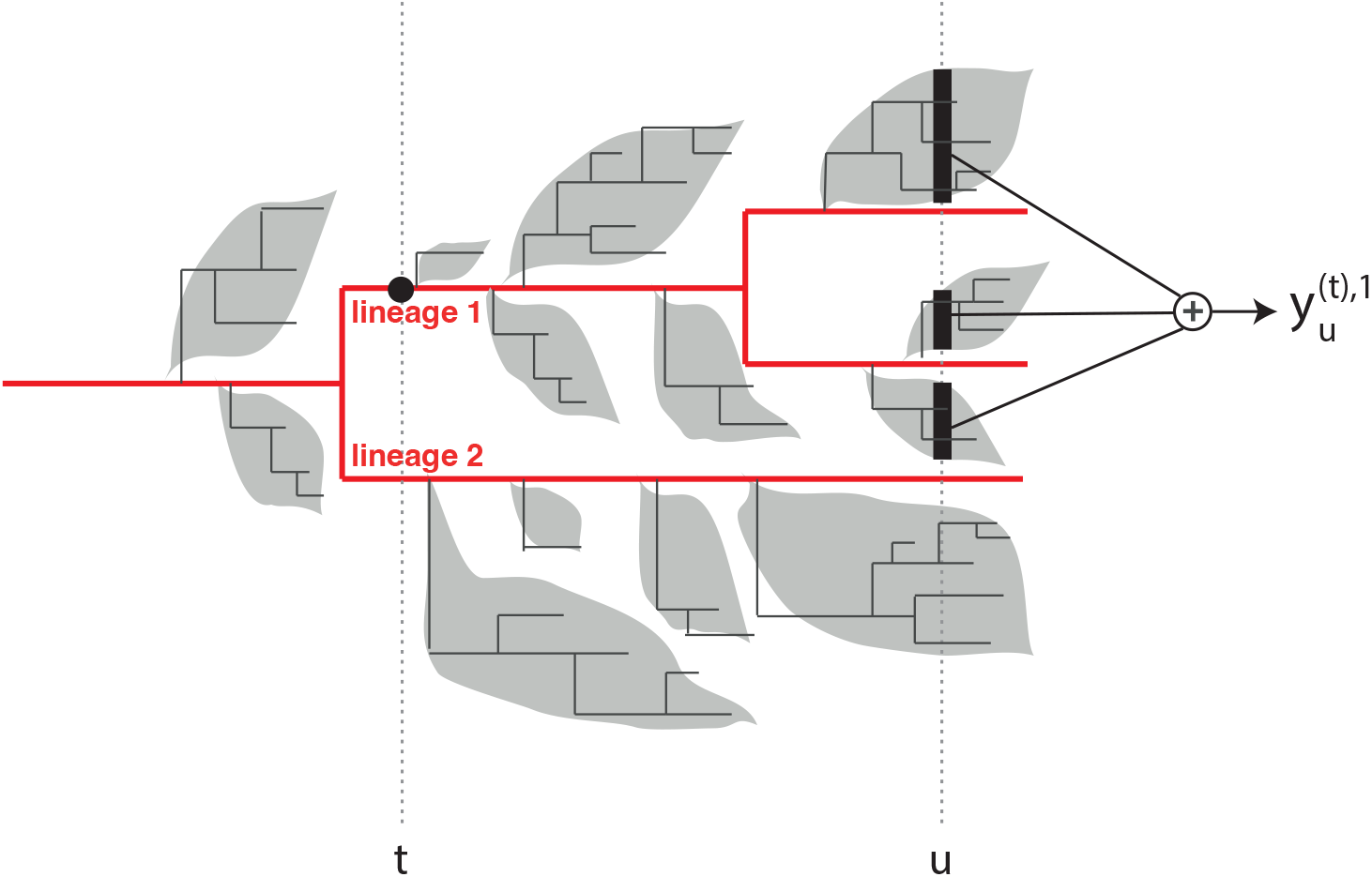
The Yule process is depicted in red. Mass is *emigrated* from the lineages, represented as grey bubbles. At the discrete level, these “mass” processes correspond to the contributions of subcritical branching processes grafted onto the immortal lineages. For each *t* ≥ 0, the total population *y*_*t*_ is obtained by summing the contributions of all the grafted sub-populations. Finally, for a fixed time *t*, the quantity 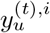 denotes the contribution at time *u* of the *i*^th^ immortal particle alive at time *t*. In the illustration, this is represented by the sum of all black intervals at time *u*.

In the following, we won’t need to go into the details of the ancestral construction of *y*, but we will only rely on a few facts that we now briefly recall. Let *L*_*t*_ be the number of immortal lineages present at time *t* (that is the number of lineages present at time *t* in the Yule process *Y*). For *i*^*th*^ immortal lineage present at time *t*, and for *u* ≥ *t*, we let 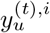 be the mass emigrated from this lineage at time *u*. See again Fig. 10 for a discrete visualization.

By the Markov property, the time shifted process 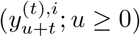 is identical in distribution to the original process *y*. According to (3.8), this implies the existence of a standard exponential random variable ℰ^(*t*),*i*^ such that

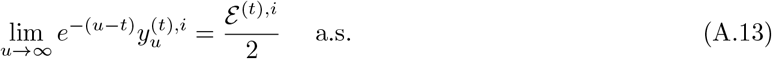

Further, conditional on *L*_*t*_, the random variables ℰ^(*i*),*t*^ are independent. Finally, we note that (A.11) and (A.13) imply the identity

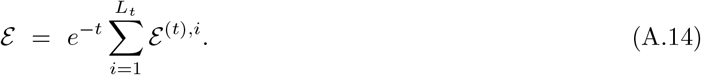

##### Step 4

Having those facts at hand, we are now ready to compute the expeactation in (A.12). We consider the expected value

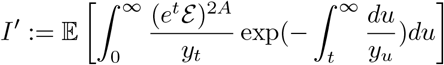

Let us denote by *T*_*c*_ the coalescing time of two lineages sampled at ∞. Since two lineages coalesce at a rate which is inversely proportional to the inverse population size, the previous quantity can be interpreted as

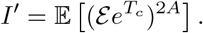

(Formally, and as in noted in Lemma 4.5. in (Etheridge *et al*., 2006), this is a variant of Perkin’s desintegration result.) From (Etheridge *et al*., 2006), 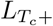 has the following distribution

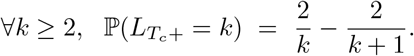

Decomposing the expected value *I*^′^ along the values for the number of immortal lineages right before coalescence, we have

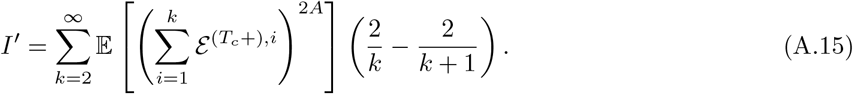

From the previous discussion, the sum 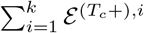 is distributed as a Gamma(1, *k*). This yields

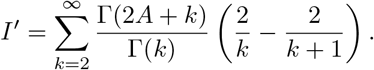

Next, a straightforward computation using properties of the Gamma functions shows that

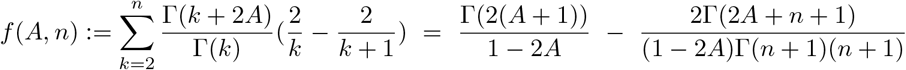

where the second term on the right hand side goes to 0 for *n* → ∞. The result then follows after combining the previous steps.

### A.3 Comparison with (Barton, 1998)

In this section, we relate the previous formula to an approach in (Barton, 1998). We briefly recall his prediction. First, consider the second-order linear ODE:

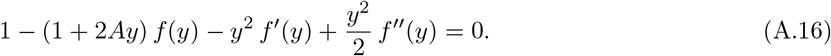

In the words of (Barton, 1998), let *h* be the solution determined by (i) the solution behaves asymptotically as *h*(*A*)*/y*^2*A*^ for large *y*, and (ii) *f* (0) = 1. According to (Barton, 1998),

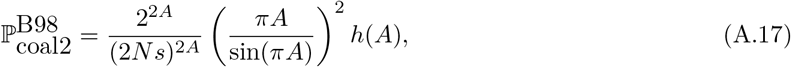

We will show that is almost consistent with the diffusion asymptotics in Eq. (A.10).

The first observation (made by D. Weissman, private communication) is that the general solution of (A.16) is

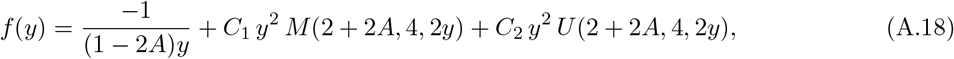

where *M* and *U* are the Kummer and Tricomi confluent hypergeometric functions respectively, and 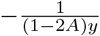 is a particular solution.

Let us now impose two boundary conditions to fix the constants *C*_1_ and *C*_2_ uniquely:

- **No exponential divergence** as *y* → ∞: since *M* (*a, b*, 2*y*) ∼ *e*^2*y*^ for large *y*, we must set *C*_1_ = 0.
- **Continuity at** *y* = 0: the particular solution 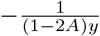 and the *y*^2^*U* term both have a 1*/y* singularity. Requiring cancellation fixes 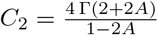.

The unique solution satisfying both conditions is:

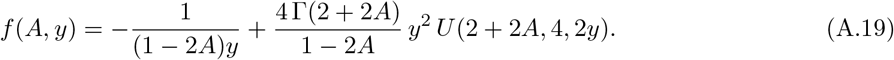

#### Behaviour at the origin

Using the expansion of *U* (*a*, 4, *z*) as *z* → 0^+^:

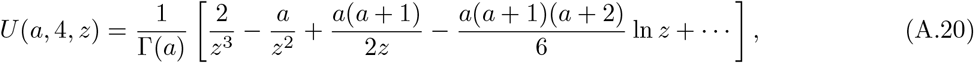

the 1*/y* singularities cancel exactly, and:

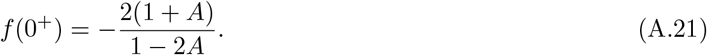

We then observe that the condition *f* (0+) = 1 imposed in (Barton, 1998) is not entirely correct. It needs to be relaxed by only imposing that *f* is continuous at 0.

**Asymptotics as** *y* → ∞

Using *U* (*a*, 4, 2*y*) ∼ (2*y*)^−*a*^ for large *y*, the two leading terms are:

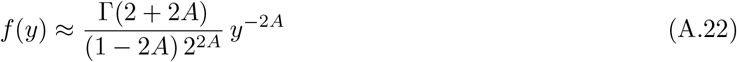

Plugging in (A.17), this yields,

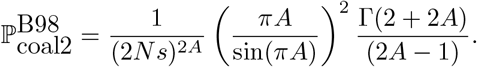

This expression coincides with (A.10) obtained from the diffusion approximation.

### A.4 Finte size correction in the diffusion approximation

#### Cut-off

The infinite sum in (A.15) is obtained by partitioning the probability space on the events 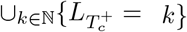. As noticed in (Durrett and Schweinsberg, 2004; Etheridge *et al*., 2006), it is then natural to introduce a cut-off in the sum (A.15) at *k* = [2*Ns*]. This leads to the definition

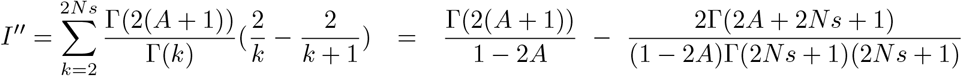

When *Ns >>* 1, a direct application of Stirling’s formula yields

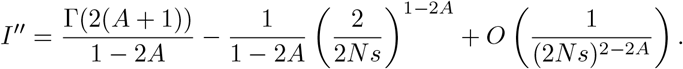

We then get the the coalescence probability

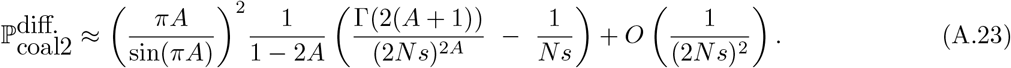

Note that compared to the first derivation (A.10), we get an extra corrective term of the order 1*/*2*Ns*.

Finally, we close this part by discussing the cut-off at [2*Ns*]. As we mentioned erleir, a similar cut-off was considered by (Durrett and Schweinsberg, 2004; Etheridge *et al*., 2006). Another rationale behind the cut-off is a posteriori: one can directly check that if we replace 2*Ns* by a variable *x* in the latter formula, then the latter expression is an asymptotic solution to the ODE A.16 BARTON (1998) as *y* → ∞.

**Transition at** *A* = 1*/*2. For *A >* 1*/*2, the exponent 1 − 2*A <* 0 so that 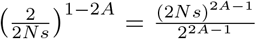 grows with *N*, and the bracket in Eq. (A.23) is dominated by the second term. The leading order asymptotics of Eq. (A.23) for *A >* 1*/*2 is therefore:

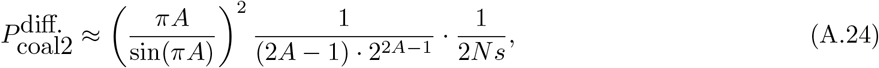

which scales as 1*/N*, consistent with the drift regime. To check that this is a continuous transition, we note that at *A* = 1*/*2 the two terms in the bracket of Eq. (A.23) cancel at leading order, and a first-order expansion yields:

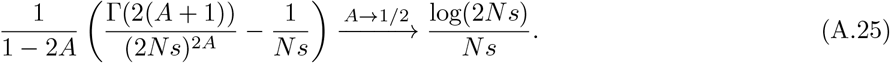

Plugging back into Eq. (A.23) yields:

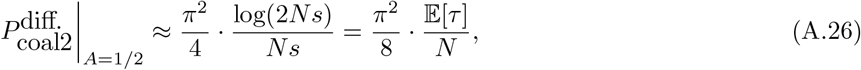

Eq. (A.26) can be further interpreted in terms of the fixation time during a sweep. By noting that 2 log(2*Ns*)*/s* ∼ E[*τ*] is the mean fixation time of the sweep (eq. (3.11)), we have:

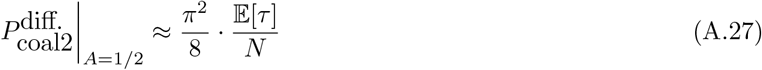

which is the neutral drift coalescence probability 1*/N* multiplied by the duration of the sweep. This indicates that at *A* = 1*/*2, the system transitions from a draft regime to a drift regime: a regime we call the dr**AI**ft regime.

## Footnotes

∗ On a more formal level, the previous observation can be made rigorous by proving that 0 is an entrance boundary (Etheridge, 2011) for the process *Z*_*t*_, i.e., there exists a solution starting at 0 and remaining strictly positive afterwards.

† Analogously to our discussion on the conditioned Wright-Fisher diffusion in Section 3.1.1 and the distinction between *Z* and 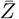 the unconditioned Feller diffusion of the 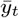 is given by: 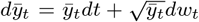. As for the Wright Fisher diffusion, the conditioning has the effect of changing the first term 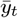 into *y*_*t*_ coth(*y*_*t*_).

‡ Note that in order to match (Barton, 1998), we replace 2*N* to *N* [diploid vs haploid] and used the identity 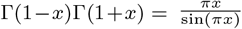 for every *x* ∈ (0, 1)

